# Hypersensitivity to distractors in Fragile X syndrome from loss of modulation of cortical VIP interneurons

**DOI:** 10.1101/2023.01.03.522654

**Authors:** Noorhan Rahmatullah, Lauren M. Schmitt, Lisa De Stefano, Sam Post, Jessica Robledo, Gunvant R. Chaudhari, Ernest Pedapati, Craig A. Erickson, Carlos Portera-Cailliau, Anubhuti Goel

## Abstract

Attention deficit is one of the most prominent and disabling symptoms in Fragile X Syndrome (FXS). Hypersensitivity to sensory stimuli contributes to attention difficulties by overwhelming and/or distracting affected individuals, which disrupts activities of daily living at home and learning at school. We find that auditory or visual distractors selectively impair visual discrimination performance in both humans and mice with FXS, but not their typically developing controls. Vasoactive intestinal polypeptide (VIP) neurons were significantly modulated by incorrect responses in the post-stimulus period during early distractor trials in WT mice, consistent with their known role as ‘error’ signals. Strikingly, however, VIP cells from *Fmr1^-/-^* mice showed little modulation in error trials, and this correlated with their poor performance on the distractor task. Thus, VIP interneurons and their reduced modulatory influence on pyramidal cells, could be a potential therapeutic target for attentional difficulties in FXS.

## INTRODUCTION

Fragile X syndrome (FXS), the most common inherited form of intellectual disability, is associated with several comorbid conditions, such as epilepsy, anxiety, aggression, autism, and sensory hypersensitivity (Bailey et al., 2008). The focus of basic and translational research efforts in animal models of FXS has been placed on investigating neural mechanisms associated with these symptoms. Behaviorally speaking, however, the most consistent feature of FXS children is their persistent inattention, impulsivity, fidgetiness, and restlessness, with most individuals with FXS meeting criteria for a diagnosis of attention deficit hyperactivity disorder, particularly the inattentive type (ADD) (Sullivan et al., 2006; Grefer et al., 2016). At the same time, individuals with FXS experience prominent sensory over-responsivity (SOR), which is characterized by exaggerated responses to certain auditory, visual, olfactory, and tactile stimuli that are innocuous to neurotypical individuals (Miller et al., 1999; Cornish et al., 2004; Van der Molen et al., 2012; Sinclair et al., 2017; Rais et al., 2018). Sometimes referred to as a sensory-modulation disorder, SOR triggers maladaptive behaviors in FXS, such as avoidance, defensive responses, or distraction and inattention (consistent with co-morbid diagnosis of ADHD), which in turn contributes to learning deficits (Kogan et al., 2004; Kaufmann et al., 2017).

Remarkably, despite the prevalence and importance of attentional difficulties in FXS, it has been understudied in animal models. Its underlying neural mechanisms, and how they might interfere with learning and cognition, are largely unknown. It has long been proposed that neuropsychiatric symptoms in FXS and other neurodevelopmental conditions (NDCs) are linked to hyperexcitability and reduced GABAergic inhibition (Rubenstein and Merzenich, 2003; Contractor et al., 2015; Contractor et al., 2021). In FXS, SOR and attentional difficulties likely engage complex interactions between excitatory neurons and several interneuron subclasses, yet these circuit dynamics have not been explored in detail during behavior.

Previously, we reported that *Fmr1^-**/-**^* mice, the best-studied animal model of FXS (Dutch-Belgian Fragile X Consortium, 1994), exhibit impairments on a go/no-go visual discrimination task compared to wild type (WT) controls—a deficit that was recapitulated in humans with FXS (Goel et al., 2018). Accurate performance in such a task requires that the animal attend to task relevant information and ignore task irrelevant information (i.e., sensory distractors) (Baluch and Itti, 2011; Zhang et al., 2014). Because *Fmr1^-**/-**^* mice exhibit SOR (Chen and Toth, 2001; Rotschafer and Razak, 2013; Rotschafer and Razak, 2014; He et al., 2017), just like people with FXS, we hypothesized that they would be unable to tune-out sensory distractors, and this would negatively impact their performance on the visual task.

Thus, we set out to investigate the intersection of attentional difficulties, SOR, and perceptual decision making in FXS, and the underlying neural mechanisms, using a visual discrimination task. We show that sensory distractors selectively impaired task performance in both *Fmr1^-/-^* mice and FXS humans. Calcium imaging in V1 showed that VIP cell activity was less modulated by visual stimuli in task naive *Fmr1*^-/-^ mice than in WT controls. Moreover, in distractor trials, VIP cells were modulated by incorrect responses in WT mice in early sessions, whereas in *Fmr1*^-/-^ mice VIP cells lacked such modulation. In fact, VIP cell modulation was correlated with the speed of perceptual learning and the ability to tune-out sensory distractors.

## RESULTS

### *Fmr1*^-/-^ mice exhibit a delay in perceptual learning compared to WT controls

To investigate symptoms of ADD in FXS, we tested the effect of sensory distractors on decision-making using the same visual discrimination task with which we previously uncovered converging perceptual learning deficits in both *Fmr1^-/-^* mice and humans with FXS (Goel et al., 2018). First, we trained awake, head restrained, water-controlled young adult (2-3 months) male and female WT and *Fmr1^-/-^* mice (n= 23 for each genotype) on a go/no-go task in which mice had to discriminate between sinusoidal gratings drifting in two orthogonal directions (see *Methods*). Specifically, they had to learn to lick for a water reward for the preferred stimulus (45° orientation) but withhold licking for the non-preferred stimulus (135° orientation; **Fig. 1A**). Correct behavioral responses included hits and correct rejections (CR), while incorrect responses (errors) included misses and false alarms (FA), both of which resulted in a ‘time-out’ punishment period of 6.5 s. Task performance was determined by the discriminability index statistic *d’* (see *Methods*). As we previously reported (Goel et al., 2018), *Fmr1^-/-^* mice showed a significant delay in leaning the visual task (defined as reaching a d’>2) compared to WT controls (on average, 4.5 d for WT mice vs. 6 d for *Fmr1*^-/-^ mice; **Fig. 1B-C**).

**Figure 1:**
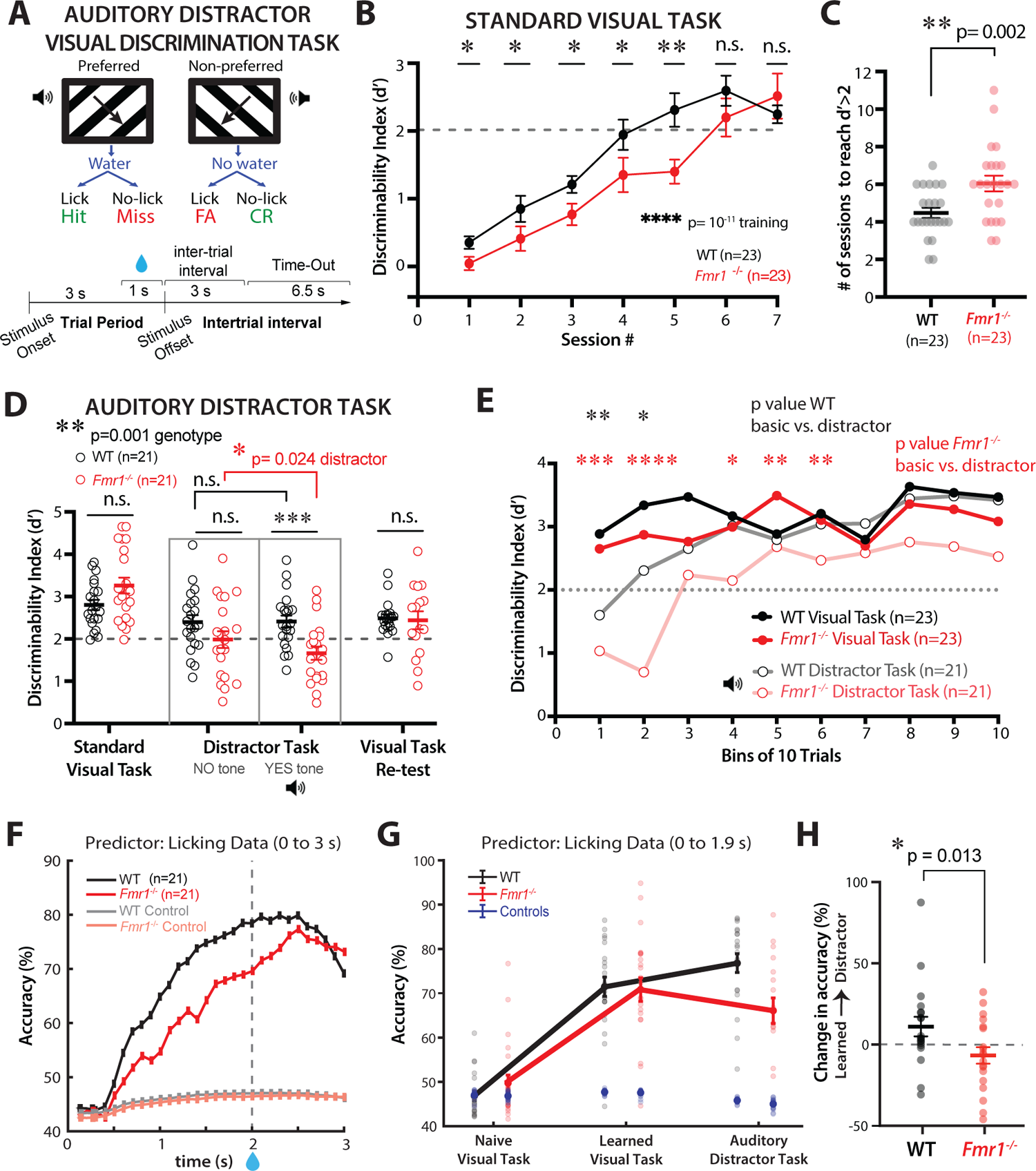
*Fmr1*^-/-^ mice exhibit a decline in performance on the visual discrimination task in the presence of sensory distractors. **A)** Illustration and timeline of behavior paradigm for auditory distractor visual discrimination task (90° degree difference between preferred and non-preferred stimuli). FA, false alarm; CR; correct rejection. Auditory distractors were presented on 50% of trials coinciding with visual stimuli. **B)** *Fmr1*^-/-^ mice exhibited delayed learning of the basic visual task (Friedman test with repeated measures for training effect, followed by Mann-Whitney test for genotype effect at each session, *F*_4,46_ = 70.15, p = 10^-11^; session 1: p = 0.05; session 2: p = 0.025; session 3: p = 0.032; session 4: p = 0.05; session 5: p = 0.007; session 6: p = 0.443; session 7: p = 0.953). Performance is measured by the discriminability index (d’). The dashed line at d’ = 2 indicates expert performance threshold. **C)** *Fmr1*^-/-^ mice took longer to achieve d’ > 2 (4.5 ± 0.3 sessions for WT mice vs. 6.0 ± 0.4 sessions for *Fmr1*^-/-^ mice; Mann-Whitney test; p = 0.002). **D)** The performance of WT and *Fmr1*^-/-^ mice was indistinguishable once they surpassed the expert threshold (2.8 ± 0.1 for WT mice vs. 3.3 ± 0.2 for *Fmr1*^-/-^ mice; Mann-Whitney test; p = 0.101). During the distractor session, there were no genotype differences in d’ on trials without distractors (2.4 ± 0.2 for WT mice vs. 2.0 ± 0.2 for *Fmr1*^-/-^ mice; Mann-Whitney test, p = 0.109). During trials with auditory distractors, *Fmr1*^-/-^ mice performed significantly worse (d’=2.4 ± 0.1 for WT mice vs. 1.7 ± 0.2 for *Fmr1*^-/-^ mice; Mann-Whitney test, p = 0.001). One the next session with the basic visual task (no distractors), there was no difference in performance between genotypes (d’= 2.5 ± 0.1 for WT mice vs. 2.4 ± 0.2 for *Fmr1*^-/-^ mice; Mann-Whitney test, p = 0.961). **E)** Performance (d’) tracked throughout the distraction task (trials grouped into bins of 10). *Fmr1*^-/-^ mice took longer to recover to “expert” level than WT controls and never reached prior levels of performance (two-way mixed ANOVA for WT basic vs. distractor; time: *F*_7,276_ = 5.1, p = 1.0E-5; task type: *F*_1,41_ = 11.7, p = 0.001; time x task type: *F*_12,444_ = 2.1, p = 0.019; two-way mixed ANOVA for *Fmr1*^-/-^ basic vs. distractor; time: *F*_7,287_ = 4.6, p = 4.9E-5; task type: *F*_1,42_ = 1.6, p = 0.214; two-way mixed ANOVA for WT learned vs. *Fmr1*^-/-^ learned; time: *F*_8,360_ = 2.7, p = 0.006; genotype: *F*_1,44_ = 0.3, p = 0.594; time x genotype: *F*_12,526_ = 1.2, p = 0.298; two-way mixed ANOVA for WT distractor vs. *Fmr1*^-/-^ distractor; time: *F*_6,203_ = 8.9, p = 6.1E-9; genotype: *F*_1,39_ = 14, p = 5.9E-4). **F)** SVM classifier from lick data predicts stimulus type with greater accuracy in WT mice than *Fmr1*^-/-^ mice. Averages of 10,000 iterations for each mouse for each 0.1 s bin of time during the 3 s stimulus period. For controls, stimuli were randomly shuffled. **G)** Accuracy of SVM classifier at different stages of task. Symbols represent individual mice. Controls (shuffled stimuli) are shown in blue. **H)** Change in accuracy of predicting stimulus type based on licking data from the learned session to the auditory distractor task (panel E), was significantly different between genotypes (+11.1 ± 26.1% for WT mice vs. −6.7. ± 22.0% for *Fmr1*^-/-^ mice; Mann-Whitney test, p = 0.013). In panels **B-H**, horizontal bars indicate mean and error bars indicate s.e.m. n values are for mice, indicated on each plot. *p < 0.05; **p < 0.01; ***p < 0.001; ****p < 0.0001.

Beyond differences in sensory processing (Goel et al., 2018) and cognitive ability as an explanation for the perceptual learning delay of *Fmr1^-/-^* mice, we considered that impulsivity and/or distractibility could also contribute significantly, given the prominent attentional difficulties in FXS. Previously, we had reported that *Fmr1^-**/-**^* mice take longer to suppress impulsive false alarm (FA) responses (i.e., persistent licking to the non-preferred stimulus) (Goel et al., 2018). Once again, we found a significantly lower percentage of CR responses and a higher percentage of FA responses in *Fmr1^-/-^* mice compared to controls on session 4, which is when WT mice have learned the task (**Supplementary Fig. 1**). This suggests that persistent licking during non-preferred stimulus trials contributed to errors, driving low performance and delayed learning in *Fmr1*^-/-^ mice.

### Decline in visual discrimination for *Fmr1*^-/-^ mice in the presence of sensory distractors

We reasoned that the higher rate of FA responses in *Fmr1*^-/-^ mice was related to baseline SOR and distraction by task-irrelevant stimuli, rather than impulsivity (Hebert, 2015; Wheeler et al., 2016). To address this, we conducted additional experiments using sensory distractors after the mice had become experts in the basic visual discrimination task. Importantly, despite the delay in learning, *Fmr1^-/-^* mice eventually reached similar expert performance levels as WT mice (**Fig. 1B, 1D**). After achieving a d’ > 2 for two consecutive days, all mice were introduced to a distractor task that included auditory in 50% of the trials, at random, coinciding with the onset of the visual stimulus. Auditory distractors consisted of loud tones (1 beep lasting 1.5 s long, 5 kHz at ∼65 dB). In separate sessions we also used visual distractors consisting of flashing lights around the monitor (white LED lights flashing 4 times for 0.5 s each with a 0.25 s interstimulus interval; see *Methods*). We found that task performance of most WT mice was unaffected by auditory distractors (**Fig. 1D**). In contrast, although *Fmr1*^-/-^ mice, on average, performed well (d’>2) on trials without distractors, their performance was significantly reduced in the presence of distractors (**Fig. 1D**). We found a similar decline in discrimination for *Fmr1*^-/-^ mice, but not for WT controls, in the presence of visual distractors (**Supplementary Fig. 2A-B**). Importantly, when *Fmr1^-/-^* mice were re-tested the following session on the standard task without distractors, they returned to expert performance levels, indistinguishable from that of WT mice (**Fig. 1D, Supplementary Fig. 2B**). This implies that it was indeed the presence of the sensory distractors that impaired task performance for *Fmr1^-/-^* mice, rather than a perceived change in the rules of the task.

The performance of both WT and *Fmr1*^-/-^ mice on the visual discrimination task, with auditory or visual distractors, was marked by significant individual variability. Some *Fmr1*^-/-^ mice performed as well as, or better than, the best WT controls, while others performed quite poorly, even in trials without distractors (**Fig. 1D, Supplementary Fig. 2B**). This suggests that the negative effect of distractors on discrimination may be particularly severe and pervasive for a subset of *Fmr1*^-/-^ mice. Interestingly, the performance of *Fmr1*^-/-^ mice on the standard visual task (without distractors) was predictive of their performance on the auditory distractor task. *Fmr1*^-/-^ mice that required more sessions to learn the basic task also showed poor performance (d’<2) on a greater number of distractor trials (**Supplementary Fig. 3A**). Furthermore, during the distractor task, there was a strong correlation between the d’ of *Fmr1*^-/-^ mice on distractor trials and their d’ on no-distractor trials (**Supplementary Fig. 3B**), which suggests the deleterious effects of distractors were pervasive.

### With practice *Fmr1*^-/-^ mice can ignore sensory distractors, but only partially

We wondered whether WT mice might be transiently affected by distractors and also whether *Fmr1*^-/-^ mice eventually learn to ignore, or tune-out, the sensory distractors and improve their visual discrimination. We calculated d’ across bins of 10 trials during the distractor session and discovered that the susceptibility of *Fmr1^-/-^* mice to distractors lasted longer than for WT controls (**Fig. 1E**; **Supplementary Fig. 2C**). Although *Fmr1^-/-^* mice reached d’>2 within 30-40 trials (depending on the distractor), they were never able to reach their prior baseline level of expertise, whereas WT mice quickly reached a d’>2 within 20 trials and their baseline performance within 30 trials. Thus, on average, *Fmr1^-/-^* mice took significantly more trials than WT mice to reach a d’>2 on the distractor tasks, or to maintain a d’>2 for two consecutive bins (**Supplementary Fig. 4A-B**). When we compared d’ between different sessions (naïve session #1, the ‘learned’ session with d’>2, and the distractor session, the only genotype difference was the lower d’ in the distractor session for *Fmr1^-/-^* mice (**Supplementary Fig. 4C**). This was associated by a higher proportion of FA responses (and fewer CRs) in *Fmr1*^-/-^ mice compared to WT controls in the presence of distractors (**Supplementary Fig. 4D**).

Licking profiles during the distractor task clearly revealed the higher proportion of FA responses observed in *Fmr1*^-/-^ mice. Whereas WT mice could suppress licking on the non-preferred trials, *Fmr1*^-/-^ mice could not (**Supplementary Fig. 5A**). The lick probability (see *Methods*) for WT mice increased early, on preferred trials (in anticipation of the water reward) but remained flat on non-preferred trials. In contrast, lick probability for *Fmr1*^-/-^ mice increased on both preferred and non-preferred trials (**Supplementary Fig. 5B**). Thus, at the time of reward, the difference in lick probability between preferred vs. non-preferred trials was larger in WT than in *Fmr1*^-/-^ mice, reflecting the greater difficulty in discriminating visual stimuli.

To demonstrate that mouse licking profiles are valid measures of performance and that they represent differences in performance between genotypes, we used the Support Vector Machine (SVM) classifier (see *Methods*) to predict the stimulus type (preferred or non-preferred) on auditory distractor trials based on licking profiles. The classifier performed with higher accuracy for data from WT than *Fmr1*^-/-^ mice, and the accuracy increased sooner after stimulus onset (**Fig. 1F**). When comparing the different sessions (naïve, learned, and distractor), the SVM classifier could more accurately predict the stimulus type on the learned session compared to the naïve session, as expected (**Fig. 1G**). However, while the accuracy was even higher for WT mice in the distractor session, it declined in *Fmr1^-/-^* mice (**Fig. 1G-H**). Altogether, these data reflect the unique and sustained susceptibility of *Fmr1*^-/-^ mice to distractors and their reduced ability to modify their behavioral responses accordingly.

### No sex differences in behavioral phenotypes of *Fmr1*^-/-^ mice

Our large sample size allowed us to look for sex differences in performance. We did not identify any sex differences for either WT or *Fmr1*^-/-^ mice as far as performance in the initial visual task or the re-test (**Supplementary Fig. 6**). Similarly, no sex differences were observed in the d’ of *Fmr1*^-/-^ mice on the distractor task. Intriguingly, d’ was significantly lower in in WT females compared to WT males in the presence of auditory distractors. (**Supplementary Fig. 6**).

### Visual discrimination performance in the presence of sensory distractors is impaired in humans with FXS

We previously demonstrated a compelling alignment of visual discrimination deficits in both humans with FXS and *Fmr1^-/-^* mice using an analogous visual discrimination task (Goel et al., 2018). To assess the translational relevance of the effects of sensory distractors in *Fmr1^-/-^* mice (and, by extension, the associated circuit dysfunction), we applied the same distractor paradigm to human subjects, with only minor modifications to make it suitable for individuals with FXS (**Fig. 2A**; see *Methods*). We administered a two-part visual task to FXS participants and age- and sex-matched typically developing controls (TDC) (n= 23 and 22, respectively; Supplementary Table 1) in a single session (*see Methods*). All of the typically developing controls (TDC males only; 20/20) achieved expert status (d’ > 2) within the first 50 trials of the standard task (without distractors) and, on average, their performance did not decline with auditory distractors (**Fig. 2B**).

**Figure 2:**
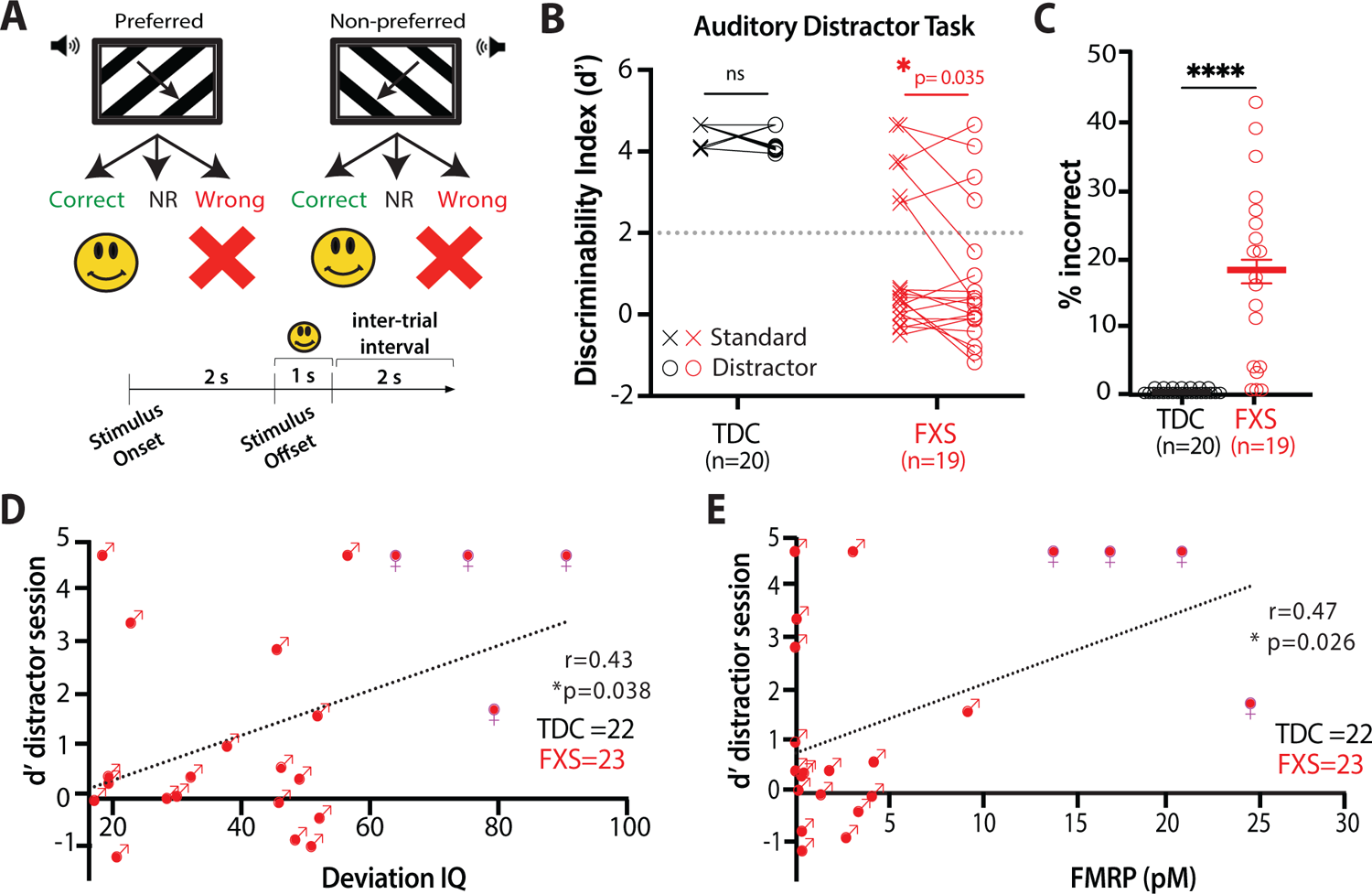
Humans with FXS exhibit a significant decline in visual discrimination with auditory distractors. **A)** Illustration and timeline of behavior paradigm for auditory distractor visual discrimination task in humans. NR, no response. Auditory distractors comprised of tones and were presented on 50% of trials. **B)** Compared to typically developing controls (TDC), FXS participants showed a significant decrease in d’ in the presence of distractors (TDC males: Standard Task d’= 4.6 ± 0.2; Distractor Task d’= 4.4 ± 0.3; p = 0.07; FXS males: Standard Task d’= 1.3 ± 0.4; Distractor Task d’= 0.8 ± 0.4; p = 0.03; paired t-test). **C)** FXS participants showed a higher percentage of incorrect responses on the distractor task (0.3 ± 0.1% in TDC vs. 17.4 ± 3.1% in FXS; t-test, p = 4.0E-5). Horizontal bars indicate s.e.m. **D)** Higher deviation IQ related to better performance in presence of the auditory distractor (r = 0.43, p = 0.038). **E)** Increased FMRP expression related to better performance with distractors (r = 0.47, p = 0.026).

Although only a subset of male FXS participants (6/19) achieved expert status (d’ > 2) within 50 trials of the standard task, on average distractors had a negative impact on the performance of participants with FXS, such that their d’ was significantly lower than in the trials without distractors (**Fig. 2B**), and this was evident in the significantly higher percentage of errors in the FXS group (**Fig. 2C**). The sensitivity to distractors in FXS participants persisted throughout the entire distractor session (not shown). The human paradigm was limited to 50 trials to maintain engagement and compliance; but it is possible that with additional trials we could have seen improved performance in FXS participants like we observed in mice. Further, in contrast to *Frm1^-/-^* mice, only male FXS participants exhibited sensitivity to the distractors.

While the performance of FXS subjects on the distractor task was on average very poor, effects were variable. Many FXS male subjects (13/19) did much worse with distractors, though a few (6/19) did surprisingly better, perhaps because they could tune out distractors and/or simply required a few more trials to learn the basic task. Thus, we calculated the absolute change in d’ triggered by auditory distractors. Compared to TD controls, the mean change in d’ was significantly larger in the presence of distracting tones for FXS participants than controls (change in d’: 273 ± 400% for FXS vs. 5 ± 6% in TDC; p=0.04). Taken together our results demonstrate that FXS male participants exhibit a similar sensitivity to sensory distractors as *Fmr1^-/-^* mice and establish this distractor assay as a useful tool with which to determine the neural mechanisms of distractibility and ADD in mouse models of NDCs.

### Deviation IQ and FMRP levels predict task performance in FXS subjects

One might expect that performance of FXS participants on the distractor task was determined by certain FXS-relevant characteristics, such as age, the degree of intellectual disability, or the expression levels of FMRP. Of note, 4 out of the 23 subjects were female (see *Methods*) and were included in the analysis examining effects of IQ and FMRP levels. We found no relationship between performance and age (|r|’s<0.32, p’s>0.14). Higher IQ was significantly correlated with better performance in presence of the auditory distractors (**Fig. 2D**; r=0.43, p=0.038), and this relationship only approached significance in the absence of the distractor (r=0.35, p=0.10; not shown). In addition, increased FMRP expression also correlated with better performance on the distractor task (**Fig. 2E**; r=0.47, p= 0.026), but not without distractors (r=0.26, p=0.24; not shown). In fact, those whose performance deteriorated the most in the presence of distractors (raw change) had the lowest FMRP expression (r=0.44, p=0.04; not shown). Interestingly, a relationship between increased caregiver-report of SOR and worsened performance in presence of auditory distractors approached significance (r = −0.42, p=0.08; not shown). Additionally, number of correct responses during distractor task related to number of correct responses during Distractibility task (r=0.61, p=0.03; not shown), and Go/No inhibitory control task (r=0.54, p=0.06). However, caregiver-reported symptoms of hyperactivity did not relate to performance (|r|’s<0.28, p’s>0.25).

### Lower selectivity of pyramidal cells in V1 of *Fmr1*^-/-^ mice

To investigate the circuit mechanisms underlying the effects of sensory distractors on visual discrimination in *Fmr1*^-/-^ mice we performed in vivo calcium imaging of pyramidal neurons in V1 with viral expression of GCaMPs, during the distractor task (**Fig. 3A-B**). Similar to the poor orientation tuning of pyramidal cells in V1 from *Fmr1^-/-^* mice (Goel et al., 2018), we observed that pyramidal cells of *Fmr1^-/-^* mice were less selective during the distractor task. This was reflected by a significantly higher percentage of them responding to both preferred and non-preferred stimuli (see *Methods*) compared to WT mice (**Fig. 3C-D**). We quantified the selectivity by calculating the single neuron performance using a receiver operating characteristic (ROC) analysis (see *Methods*). The area under the curve for pyramidal neurons in *Fmr1^-/-^* mice was significantly smaller than for WT mice (**Fig. 3E-F**), indicating that the fraction of distractor trials correctly discriminated by pyramidal neuron firing was lower in *Fmr1^-/-^* mice, which likely reflects their lower selectivity (broader tuning), thereby impairing behavioral performance on the distractor task.

**Figure 3:**
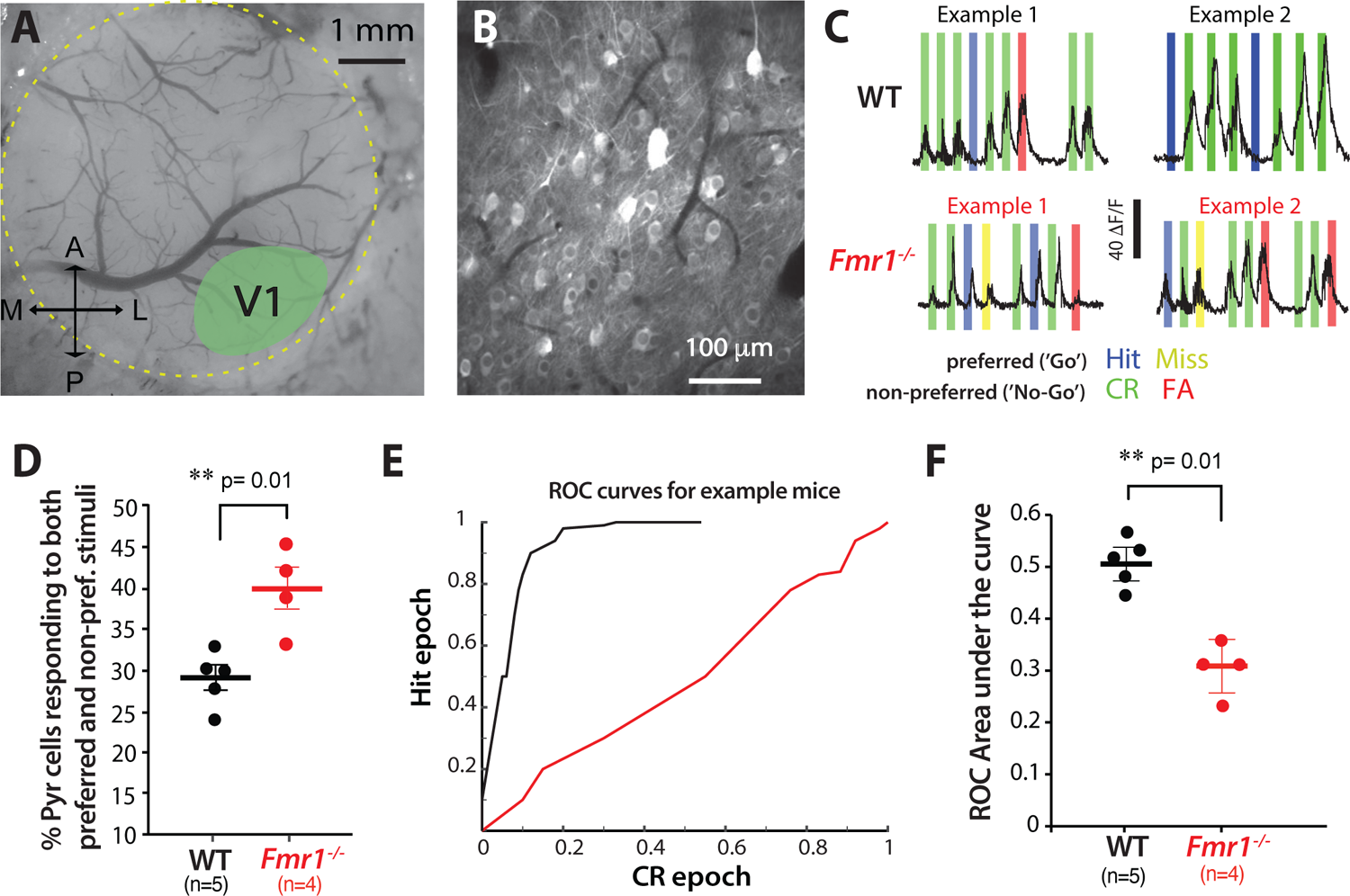
Reduced orientation selectivity of pyramidal cells after learning in *Fmr1*^-/-^ mice. **A)** Example cranial window showing approximate location of V1(A: anterior, P: posterior, M: medial, and L: lateral. **B)** Representative field of view for in vivo two-photon calcium imaging of layer 2/3 pyramidal (Pyr) neurons in V1 expressing GCaMP7f. **C)** Traces of visually evoked calcium transients for two example Pyr neurons during the distractor task from WT and *Fmr1*^-/-^ mice. Responses (as determined by changes in jGCaMP7f fluorescence intensity, ΔF/F) were seen across both preferred and non-preferred trials in *Fmr1*^-/-^ mice, whereas Pyr neurons in WT mice were tuned to either or. **D)** The percentage of Pyr cells that respond to both preferred and non-preferred stimuli on the distractor task was higher in *Fmr1*^-/-^ mice (28.7 ± 1.4% for WT mice vs. 39.6 ± 2.6% for *Fmr1*^-/-^ mice; t-test, p = 0.003). **E)** Example ROC curves (*see Methods*) for the data shown in example #2 in panel C **F)** The mean area under the curve for the ROC was smaller in *Fmr1^-/-^* mice, consistent with the reduced selectivity of Pyr cells during the distractor task (0.5 ± 0.02 for WT mice vs. 0.3 ± 0.03 for *Fmr1*^-/-^ mice; t-test, p = 0.0002).

### Reduced modulation of VIP interneurons in V1 of *Fmr1*^-/-^ mice by visual stimuli

The distractor task requires mice to ignore task irrelevant stimuli (i.e., the sensory distractors) to maintain performance and receive a reward. However, *Fmr1^-/-^* mice exhibited higher rates of FA responses than WT mice and a decreased ability to adapt to the changing conditions of the task. Based on the differences in pyramidal cell selectivity, we hypothesized that reduced modulation of VIP neurons in *Fmr1*^-/-^ mice might also contribute to impaired visual discrimination, especially in the presence of distractors. Indeed, VIP neurons serve as a principal circuit mechanism to increase the gain of pyramidal ensembles (Pi et al., 2013; Hangya et al., 2015; Turi et al., 2019) thereby contributing to novelty detection (Garrett et al., 2020).

We used in vivo 2-photon calcium imaging to simultaneously record the activity of VIP and pyramidal neurons in V1 of WT and *Fmr1*^-/-^ mice (n= 13 and 12, respectively). During the cranial window surgery, we injected a Cre-dependent virus into V1 of VIP-Cre mice x Ai9;TdTomato mice (see *Methods*) to selectively express TdTomato and jGCaMP7f in VIP cells, in addition to expressing GCaMP7f in excitatory neurons (**Fig. 4A-C**). Initially, we recorded activity during passive visual stimulation in task naïve mice. We discovered pronounced genotype differences in visually evoked activity of VIP cells (**Fig. 4D**). VIP neurons in WT mice showed prominent but non-selective visually evoked responses to sinusoidal gratings drifting in different directions, characteristic of intra-population coupling of this cell type (Karnani et al., 2016). In stark contrast, VIP cells in *Fmr1^-/-^* mice exhibited minimal modulation by visual stimuli. Instead, we observed persistently elevated activity with slow fluctuations that did not correspond to individual stimulus epochs (**Fig. 4D**). The mean magnitude of visually evoked activity of VIP cells was significantly higher in *Fmr1^-/-^* mice than in WT mice (**Fig. 4E**), while the fraction of stimulus-responsive VIP cells was significantly lower in *Fmr1^-/-^* mice (**Fig. 4F**). We did not find differences in the total number of active VIP cells imaged per field of view between WT and *Fmr1^-/-^* mice (**Supplementary Fig. 7**). To further determine the extent to which VIP cells were responsive to visual stimuli (drifting gratings), we calculated a modulation index (see *Methods*) and found that VIP cells from *Fmr1^-/-^* mice were significantly less modulated by visual stimulation than WT mice (**Fig. 4G**). In fact, only 57.3% of VIP cells showed any significant modulation by visual stimuli in *Fmr1^-/-^* mice, compared to 73.2% in WT mice (**Fig. 4H**).

**Figure 4:**
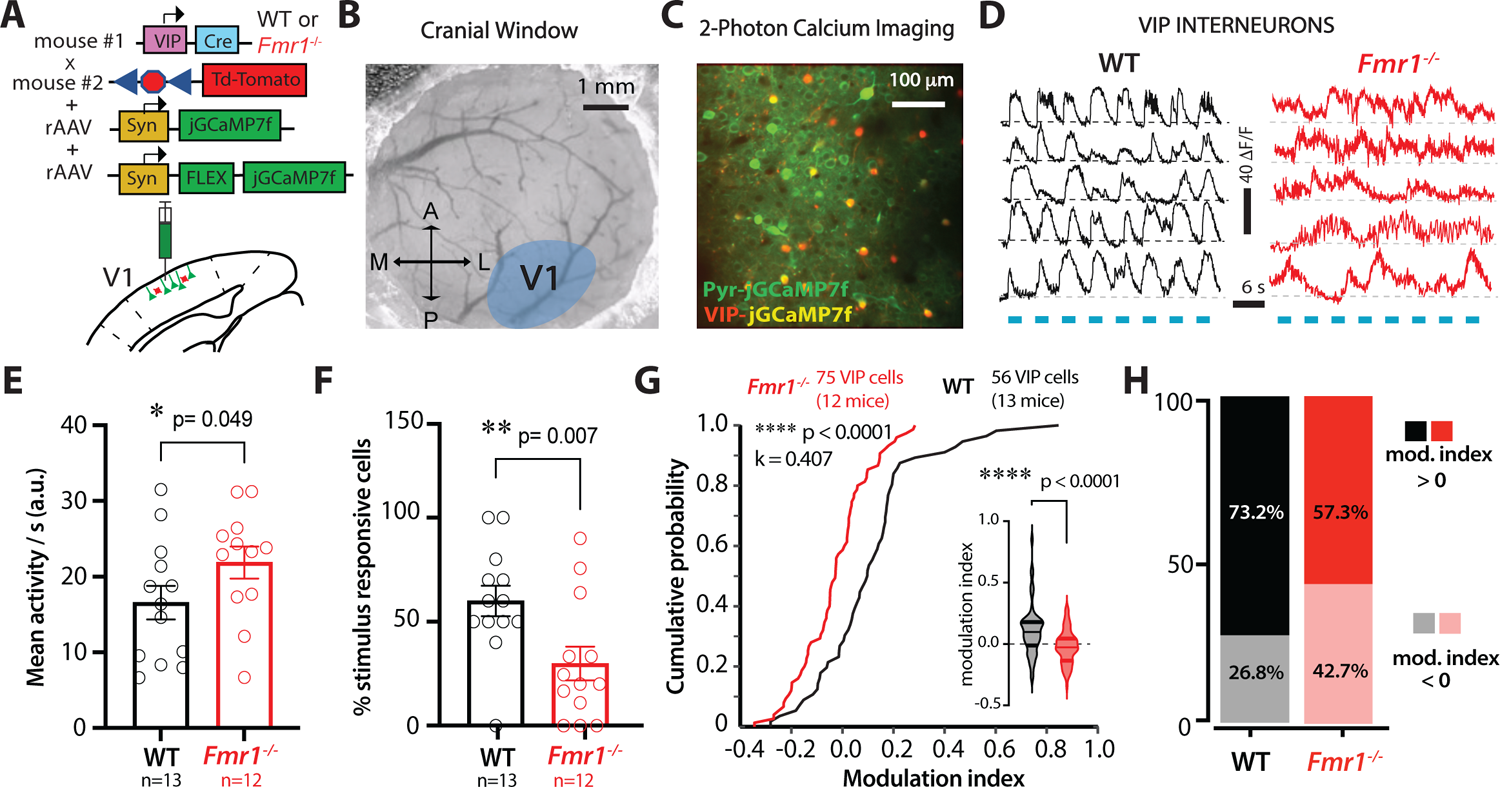
Reduced modulation by visual stimuli of VIP activity in *Fmr1*^-/-^ mice. **A)** Illustration of co-injection of rAAV-syn-jGCaMP7f and rAAV-syn-FLEX-jGCaMP7f in V1 in VIP-cre x ai9 mice (td-Tom). **B)** Example cranial window over V1 (labels as in Fig. 3A). **C)** Representative field of view for in vivo two-photon calcium imaging of Pyr (green) and VIP neurons (red-yellow). **D)** Example traces of visually evoked calcium transients for VIP neurons in WT and *Fmr1*^-/-^ mice. Blue bars represent epochs of sinusoidal gratings (drifting in 8 directions). **E)** Mean visually evoked activity of VIP cells (as measured by the area under the trace; a.u.) was higher in *Fmr1*^-/-^ mice (16.6 ± 2.2 for WT mice vs. 21.9 ± 2.1 for *Fmr1*^-/-^ mice; Mann-Whitney test, p = 0.049, Cohen’s *d* = 0.691). Symbols denote individual mice. **F)** There were fewer VIP neurons in *Fmr1*^-/-^ mice that were responsive to the visual stimuli (60 ± 7.3% for WT mice vs. 29.9 ± 8.1% for *Fmr1*^-/-^ mice; Mann-Whitney test, p = 0.007, Cohen’s *d* = 1.084). **G)** A cumulative probability plot showing reduced VIP cell modulation by visual stimuli for *Fmr1*^-/-^ mice as measured by a modulation index (two-sample Kolmogorov-Smirnov test, p = 2.81E-05, k = 0.407). Violin plot inset shows reduced VIP modulation in *Fmr1*^-/-^ mice (values indicate change in activity as a result of visual stimuli compared to gray screen (+0.11 ± 0.03 for WT mice vs. −0.03 ± 0.02 for *Fmr1*^-/-^ mice; Mann-Whitney test, p = 8.44E-6 Cohen’s *d* = 0.783). **H)** The percentage of VIP cells that were positively modulated by the visual stimuli was smaller in *Fmr1*^-/-^ mice (57.3%) than in WT controls (73.2%).

### VIP interneurons fail to signal incorrect responses in *Fmr1*^-/-^ mice

We next examined whether this reduced dynamic range of VIP neurons persists during the distractor session, which could prevent *Fmr1*^-/-^ mice from using error signals to adjust decisions. When mice perform sensory discrimination tasks, cortical VIP neurons are more active during incorrect responses, which suggests they function as error signals, providing reinforcement feedback that is important for learning (Pi et al., 2013). Considering the increase in incorrect responses in the distractor session, we next recorded VIP activity in the presence of distractors. We found that during auditory distractor trials, VIP neurons in WT mice (n= 42) responded to both preferred and non-preferred visual stimuli; however, their mean activity during the stimulus period was greater on error trials (Misses, FAs) than on correct trials (Hits, CRs) (see example cells in 1 mouse in **Fig. 5A**). Interestingly, VIP cells in WT mice showed persistently elevated activity during such error trials well beyond the stimulus period (**Fig. 5A**), consistent with previous reports (Pi et al., 2013). In contrast, VIP cells from *Fmr1^-/-^* mice (n= 44) seemed to be much less modulated by visual stimulation during the distractor task (see example in **Fig. 5B**), just as we saw in task-naïve mice (**Fig. 4G-H**).

**Figure 5:**
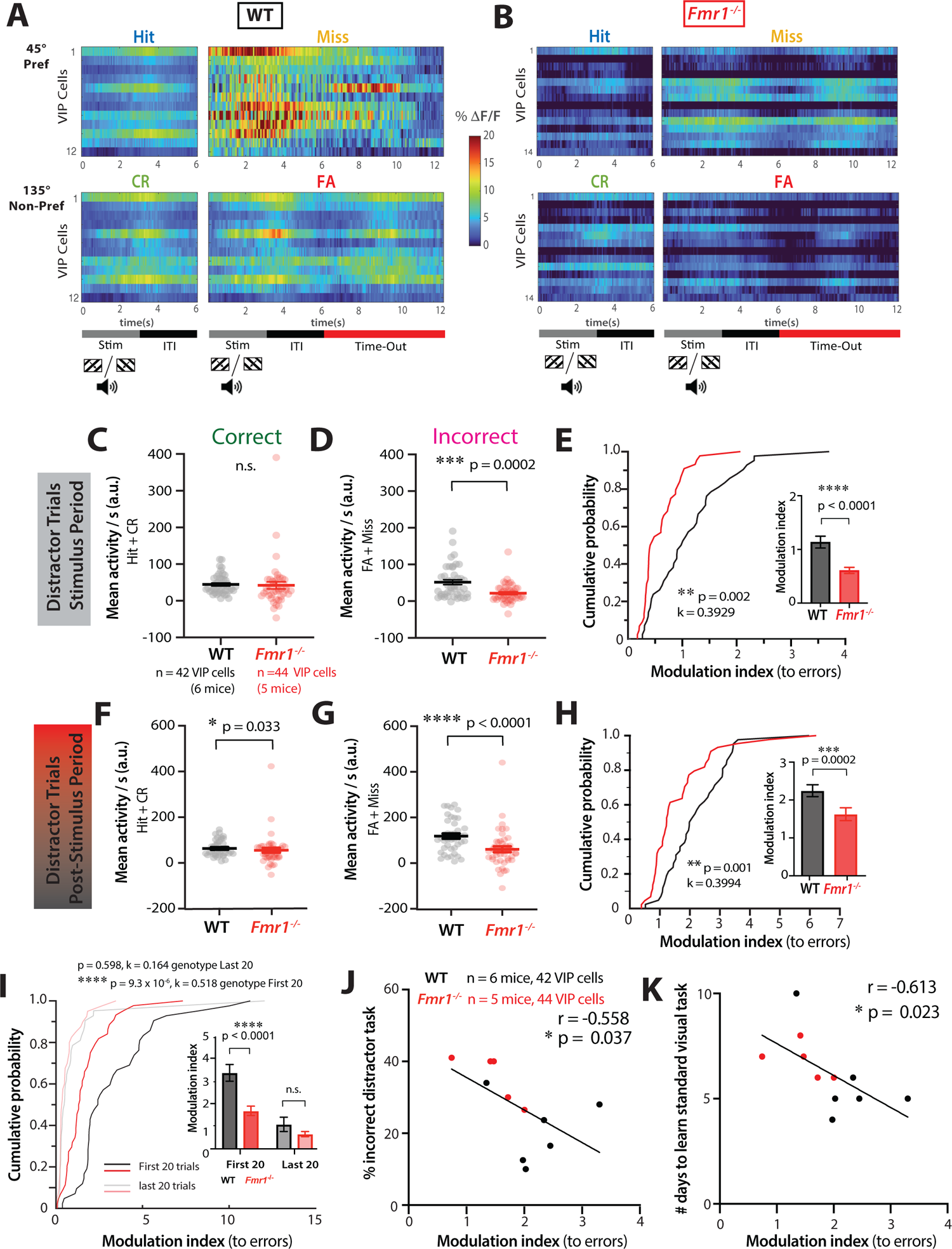
Reduced modulation of VIP activity by incorrect responses in *Fmr1*^-/-^ mice correlates with delayed learning and poor performance on the distractor task. **A)** Rasters of individual VIP neuron activity in an example WT mouse sorted by trials of different response type – hits, misses, CRs, FAs. Two-photon calcium imaging was performed during the distractor task and recordings were made from VIP neurons. Each row represents an average across all trials of the specific response type for that neuron. The timeline at the bottom denotes the visual stimulus presentation (0-3 s), the intertrial interval (3-6 s), and the time-out (6-12.5 s, only for misses and FAs). **B)** Same as panel A but for VIP neurons from an example *Fmr1*^-/-^ mouse. Note the lack of modulation of VIP cell activity during the stimulus period or the post-stim. Period **C)** Mean VIP activity during the stimulus period on auditory distractor trials (as measured by the area under the trace per s) was similar across genotypes during correct trials (hits and CRs) (44.6 ± 27.1 for WT vs. 42.1 ± 65.3 for *Fmr1*^-/-^; Mann-Whitney test, p = 0.055). **D)** Mean VIP activity during error trials (FAs and misses) was significantly lower in *Fmr1*^-/-^ mice (51.7 ± 44.5 for WT vs. 21.8 ± 25.7 for *Fmr1*^-/-^; Mann-Whitney test, p = 2.0E-4, Cohen’s *d* = 0.823). **E)** VIP cell modulation by incorrect responses (errors) was significantly lower in *Fmr1*^-/-^ mice (two-sample Kolmogorov-Smirnov test, p = 0.002, k = 0.3929). Bar graph inset showing VIP cell modulation by errors (1.1 ± 0.7 for WT vs. 0.6 ± 0.4 for *Fmr1*^-/-^; Mann-Whitney test, p = 2.0E-5, Cohen’s *d* = 0.916). **F)** Mean VIP activity was slightly lower in *Fmr1*^-/-^ mice for correct responses during the post-stimulus period on distractor trials (59.6 ± 34.2 for WT vs. 50.4 ± 68.0 for *Fmr1*^-/-^; Mann-Whitney test, p = 0.033, Cohen’s *d* = 0.17). **G)** Mean VIP activity during error trials was much lower in *Fmr1*^-/-^ mice(123.6 ± 72.4 for WT vs. 65.3 ± 85.9 for *Fmr1*^-/-^; Mann-Whitney test, p = 4.8E-5, Cohen’s *d* = 0.734). **H)** VIP cell modulation by errors was significantly reduced in *Fmr1*^-/-^ mice in the post-stimulus period (two-sample Kolmogorov-Smirnov test, p = 0.001, k = 0.3994). Bar graph inset showing VIP cell modulation by errors (2.2 ± 1.0 for WT vs. 1.6 ± 1.1 for *Fmr1*^-/-^; Mann-Whitney test, p = 2.0E-4, Cohen’s *d* = 0.573). **I)** When comparing different trials across the distractor session, VIP cell modulation by errors during the post-stimulus period was highest for WT mice in the first 20 trials, and significantly lower for *Fmr1*^-/-^ mice (two-sample Kolmogorov-Smirnov test, p = 9.3E-6, k = 0.518). Bar graph inset showing VIP cells were less modulated by errors in *Fmr1*^-/-^ mice on the first 20 distractor trials (3.4 ± 2.4 for WT vs. 1.7 ± 1.3 for *Fmr1*^-/-^; Mann-Whitney test, p = 7.0E-6, Cohen’s *d* = 0.868), but there was no significant difference on the last 20 distractor trials (1.1 ± 2.1 for WT vs. 0.7 ± 0.7 for *Fmr1*^-/-^; Mann-Whitney test, p = 0.231). A mixed-effects analysis revealed a significant effect of time, effect of genotype, and interaction effect of time x genotype (three-way mixed ANOVA; time: *F*_1,80_ = 39, p = 1.9E-9; genotype: *F*_1,84_ = 14.5, p = 0.0003; time x genotype: *F*_1,80_ = 5.6, p = 0.021). **J)** The average modulation index of VIP cells during the post-stimulus period of all auditory distractor trials for individual mice was negatively correlated with the % incorrect responses on the distractor task (Pearson’s r, r = −0.558, p = 0.037). A ‘best fit’ regression line is shown. **K)** The modulation index of VIP cells was negatively correlated with number of days to learn the standard visual task (Pearson’s r, r = −0.613, p = 0.023). In panels **C-I,** horizontal bars indicate mean and error bars indicate s.e.m; n values are for mice and cells, indicated on the figure. *p < 0.05; **p < 0.01; ***p < 0.001; ****p < 0.0001.

To quantify the differences in VIP responses throughout the distractor trial period, we first examined VIP responses during the stimulus presentation epochs (**Fig. 5C-E**). On average, there was no difference between genotypes in visually evoked activity of VIP cells during correct trials (**Fig. 5C**), but VIP activity was significantly reduced in *Fmr1^-/-^* mice on error trials (**Fig. 5D**). Moreover, the modulation index for VIP cell activity in response to errors (see *Methods*) was significantly lower in *Fmr1^-/-^* mice than in WT mice (**Fig. 5E**). Similar to data shown in Supplementary Fig. 7, differences in modulation of VIP activity could not be explained by the number of VIP cells that were active in each field of view (not shown). These genotype differences VIP cell activity, including the lack of modulation by error trials (FA + Miss), were also found during the post-stimulus period (**Fig. 5F-H**). Thus, VIP cells continue to fire during the post-stimulus period in WT mice, but less so in *Fmr1^-/-^* mice. Overall, these data show that a reduced dynamic range of VIP cell activity in *Fmr1^-/-^* mice prevents error detection that is critical for stimulus discrimination during the distractor task.

To determine if this is specific to the presence of a sensory distractor, we looked more closely at trials without distractors present. We observed very similar differences in the activity of VIP cells in *Fmr1^-/-^* mice, including reduced mean firing and reduced modulation by incorrect responses (**Supplementary Fig. 8A-F**). However, the lack of modulation of VIP activity was much more pronounced in the presence of distractors (e.g., compare **Fig. 5E** to **Supplementary Fig. 8C** and **Fig. 5H** to **Supplementary Fig. 8F**). This aligns well with our behavioral observations that *Fmr1^-/-^* mice performed even worse in the distractor task when distractors were present (**Fig. 1D**).

The most significant declines in behavioral performance of both WT and *Fmr1^-/-^* mice occurred at the beginning of the distractor task (**Fig. 1E**). For WT mice this drop in d’ was transient and they returned to expert performance within 20 trials. Hence, we hypothesized that error trial modulation of VIP cell activity would be greatest during the first few distractor trials and then attenuate in later trials, at least in WT mice. We compared VIP modulation during the first 20 trials, the middle 20 trials, and the last 20 trials. As we expected, mean visually evoked activity during error trials was significantly lower in *Fmr1^-/-^* mice than in WT controls for the first 20 trials, but was similar between genotypes in the last 20 trials, throughout the stimulus and post-stimulus periods (**Supplementary Fig. 9A-D**). Moreover, the modulation index for VIP cell activity in response to errors was significantly higher for WT mice during the first 20 trials compared to the last 20 trials (**Fig. 5I; Supplementary Fig. 9E**). Importantly, the modulation index was significantly lower for *Fmr1^-/-^* mice in the first 20 trials, but not in the last 20 (**Fig. 5I**).

Altogether, these results argue that a lack of modulation of VIP activity by incorrect responses in *Fmr1^-/-^* mice (V1 is unable to tune-out sensory distractors) impairs their behavioral performance (**Fig. 1D**). In support of this argument, we found a significant negative correlation between the modulation index of VIP cells by error trials and the percentage of incorrect responses on the distractor task, such that animals with higher modulation made fewer mistakes (**Fig. 5J; Supplementary Fig. 9F**). Moreover, we find a similar relationship between VIP modulation index and the number of days it takes mice to learn the standard visual discrimination task (**Fig. 5K; Supplementary Fig. 9G**), which implies that VIP modulation is important for learning. The correlations were particularly strong for VIP activity during the post-stimulus period, which is presumably when the animal recognizes the incorrect outcome. Our findings suggest that the reduced modulation of VIP cells in *Fmr1^-/-^* mice indicates a failure to communicate a reinforcement feedback signal following an error and impairing stimulus discrimination on the distractor task

## DISCUSSION

SOR is a prevalent symptom in neurodevelopmental conditions (NDCs) that can trigger anxiety and inattention, with dire consequences on learning and cognition. (Robertson and Baron-Cohen, 2017). We set out to investigate the impact of sensory distractors on perceptual learning and sensory discrimination, as it relates to attentional difficulties in FXS. We implemented a highly translational visual discrimination assay in *Fmr1^-/-^* mice and FXS patients and followed a symptom-to-circuit approach to identify specific circuit-level differences using in vivo two-photon calcium imaging in V1. Our findings clearly demonstrate the debilitating consequences of sensory distractors on sensory processing in both FXS humans and mice – an inability to ignore task-irrelevant tones and flashing lights. Our mouse data identifies a novel circuit mechanism for these behavioral deficits–a lack of modulation of VIP interneurons by error signals, particularly in the post-stimulus period. We believe that such a restricted dynamic range of VIP cell activity could be implicated in other NDCs characterized by SOR and inattention.

The present study accomplished two goals. The first was to provide a plausible link between SOR and learning deficits by showing how sensory distractors negatively impact performance on a visual discrimination task. A unique aspect of our approach is that we employed a parallel behavioral paradigm in humans and mice, identifying very similar deficits across species, which strengthens the face validity of our assay. The second was to shed light onto the circuit mechanisms underlying this phenomenon, and identifying VIP cell firing as a potential target for future clinical interventions.

VIP interneurons play an instrumental role in sensory cortical networks (e.g., V1), integrating inputs from other regions: 1) bottom-up sensory signals (from thalamus) with top-down inputs from higher order brain regions that help the animal select task-relevant information and suppress task-irrelevant information (i.e., tune-out distractors) (Baluch and Itti, 2011; Miller and Buschman, 2013; Zhang et al., 2014; Clark et al., 2015); 2) subcortical neuromodulation from basal forebrain improving the reliability of V1 responses (Bennett et al., 2013; Carcea and Froemke, 2013; Pi et al., 2013); 3) cortico-cortical connections resulting in activation of V1 to auditory input (Iurilli et al., 2012; Deneux et al., 2019). The reduced modulation of VIP cell activity we observed in *Fmr1^-/-^* mice during different response types on the distractor task (**Fig. 5**) suggests the animals could not learn from their mistakes. Indeed, VIP cells in primary auditory cortex (A1) are known to be recruited by reinforcement signals during an auditory discrimination task and their firing was enhanced during errors/punishment (Pi et al., 2013). This is consistent with our results in WT mice, who displayed prominent error signals in VIP cells during the first few trials of the distractor task, which led to improvement in performance. Instead, the smaller dynamic range of VIP cells in *Fmr1^-/-^* mice was particularly pronounced during the ‘time-out’ period, thereby preventing the generation of a feedback error signal in the network (**Fig. 6**). This model is supported by experiments showing how optogenetic manipulations of VIP cells enhance sensory processing (Zhang et al., 2014)

**Figure 6:**
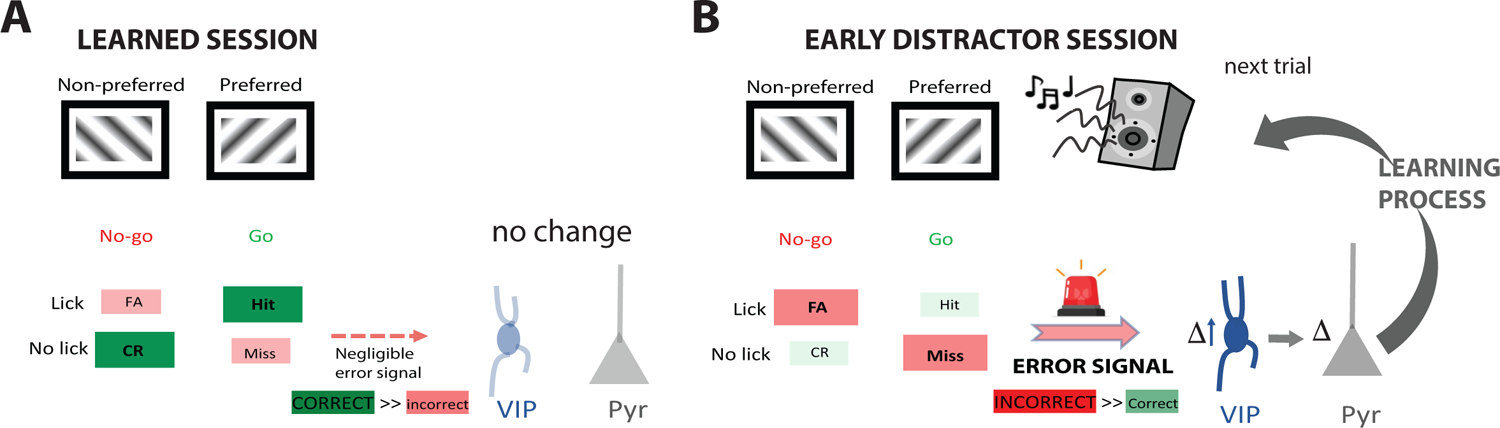
Proposed model for V1 circuit differences in *Fmr1*^-/-^ mice contributing to distractor susceptibility. Our data suggests a smaller dynamic range of VIP cell activity in V1disrupts their ability to control the gain and selectivity of their pyramidal cell partners, via disinhibition. **A)** During the learned session, when mice have become experts on the visual task, the number of correct responses is greater than errors (mice learned to increase attention to minimize FAs and Misses). In the absence of errors on the task modulation of VIP cell activity is minimal. **B)** During the initial trials on the distractor task, incorrect responses dominate, resulting in an error signal (negative reinforcement). This triggers an increase in VIP cell firing (higher VIP modulation), which presumably suppresses the effect of the distractor and leads to learning on the next trial. In contrast, VIP neurons in *Fmr1*^-/-^ mice are not modulated by error signals, contribute to distractor susceptibility, so they show reduced learning on the next trial (not shown)

Sensory-guided behavior is heavily influenced by top-down signals that filter out irrelevant sensory inputs, and enhance neuronal representation of behaviorally relevant information (Miller and Buschman, 2013; Zhang et al., 2014; Clark et al., 2015; He et al., 2017). Anterior cingulate cortex (ACC) is one area implicated in tasks involving attention, stimulus change, and error detection (Garavan et al., 2002; Zhang et al., 2014; Fiser et al., 2016). Long-range glutamatergic projections from ACC have been shown to modulate sensory processing (Baluch and Itti, 2011; Zhang et al., 2014). This process is largely mediated by ACC activation of VIP cells, which selectively enhance pyramidal cell responses through a disinhibitory process (see below) and improving discrimination (Karnani et al., 2016; Zhang et al., 2016). Interestingly, human studies focusing on attention and inhibitory control in autism and FXS have demonstrated an uncharacteristic low activation profile in ACC during attentive tasks (Chan A.S et al., 2011). Further, a recent study using *Fmr1*^-/-^ mice showed a disruption in cholinergic tone in ACC, resulting in hyperconnectivity of local ACC inputs and contributing to attention deficits (Falk et al., 2021). Follow-up studies may seek to investigate long-range inputs from the ACC to determine whether differences in top-down control contribute to the changes we observed in VIP cells of *Fmr1^-/-^* mice, as it relates to selective attention and sensory discrimination.

VIP cells form part of a cortical disinhibitory circuit that ultimately increases the gain of pyramidal cells and facilitates learning (Letzkus et al., 2011; Jiang et al., 2013; Pi et al., 2013; Fu et al., 2014; Ren et al., 2022). Activation of VIP cells suppresses SST and PV cell firing, releasing pyramidal cells from the inhibitory effects and increasing visual responses during stimulus-evoked activity (Karnani et al., 2016). Previously, we reported that hypoactivity of PV cells in *Fmr1^-/-^* mice contributes to their delayed learning of the visual task (Goel et al., 2018). Although our current results align well with those previous findings, the influence of prefrontal afferents on VIP cells in V1 could also impact other interneuron subtypes and, in turn, modulate neuronal oscillations and sensory coding (Lee et al., 2018). Future studies using sensory distractor tasks in humans while recording neural activity (visually evoked potentials, EEG) will be needed to support our findings in mice.

Our studies also demonstrate how, despite the significant group effects between genotypes, there is significant variability across individual *Fmr1^-/-^* and WT mice. Importantly though, we could correlate this variability to various metrics of neuronal activity and behavior. Overall our data highlights that a better understanding of how distracting sensory stimuli affect attention and cognitive performance, and of the underlying circuit changes in the brain, could be critical to the development of new symptomatic treatments for FXS and other NDCs. Our parallel “mouse/human” perspective, derived from a circuit-level understanding of FXS symptoms, is an exciting approach for such discoveries.

## METHODS

### Experimental animals

All experiments followed the U.S. National Institutes of Health guidelines for animal research, under animal use protocols approved by the Chancellor’s Animal Research Committee and Office for Animal Research Oversight at the University of California, Los Angeles and at the University of California, Riverside (ARC #2007-035 and ARC #2019-0036, respectively). Experiments in Fig. 1 used male and female FVB.129P2 WT mice (JAX line 004828) and *Fmr1^-/-^* mice (Dutch-Belgian Fragile X Consortium, 1994) (JAX line 004624) and experiments in Fig. 2-4 used male and female VIP-Cre mice (JAX line 010908) that were crossed to the Ai9 (td-Tomato) reporter line (JAX line 007909) and the resulting VIP-Cre x Ai9 mice were back crossed to FVB WT and *Fmr1^-/-^* mice for 8 generations. All mice were housed in a vivarium with a 12/12 h light/dark cycle, and experiments were performed during the light cycle. The FVB background was chosen because of its robust breeding, because FVB *Fmr1^-/-^* dams are less prone to cannibalizing their pups, and because FVB *Fmr1^-/-^* mice have well-documented deficits in sensory processing (Contractor et al., 2015). We used separate homozygous litters of WT and *Fmr1^-/-^* mice rather than littermate controls because littermates of different genotypes tend to receive unequal attention from the dam (Zupan and Toth, 2008), which may affect the health and behavior of *Fmr1^-/-^* pups, biasing results. To avoid issues with genetic drift, we obtained new WT and *Fmr1^-/-^* breeders from Jackson Labs at regular intervals (every 1-1.5 years).

### Go/No-go visual discrimination task for head-restrained mice

A go/no-go visual discrimination task similar to that outlined in our prior study (Goel et al., 2018) was administered to awake, head restrained young adult mice (beginning at 6-8 weeks) able to run on an air-suspended polystyrene ball treadmill. Beginning 5-7 d of recovery from head bar attachment and/or cranial window surgery, mice went through handling, habituation, and pretrials. Mice were handled gently for 5 min each day for 3 d until they were comfortable with the experimenter and would willingly transfer from one hand to the other. This was followed by water restriction, during which mice were given a rationed daily supply of water according to their weight, and the habituation phase. During habituation, mice were acclimated to the behavior rig (and microscope for mice that were imaged) for 15 min each day. They were first head-restrained and placed on the polystyrene ball and then gradually introduced to the visual stimuli, the lickport, the red light illuminating the ball, various sounds (fans circulating air in the rig, vacuum pump for water reward, scan mirrors), and objective for imaging. We started water restriction a few days before pretrials to motivate the mice to lick during pretrials (Guo et al., 2014). After habituation and achieving ∼15-20% weight loss, mice were advanced to the pretrial phase.

During pretrials, sinusoidal gratings drifting in eight different directions (temporal frequency of 2 Hz, spatial frequency of 0.01 cycles/degree and 100% contrast) were displayed on the monitor’s screen (23” display; Dell P2311HB or ThinkVision T24i-10). The monitor was placed 25 cm away from the mouse and stimuli were presented at random for 3 s, with a 3 s intertrial interval (ITI) where a grey screen was presented. Each stimulus was initially coupled with a small water reward (∼3 µL) dispensed from the lickport beginning at 2 s after the onset of stimulus presentation and up until the 3 s timepoint when the stimulus ended (water reward ‘window’). Licking by the mouse interrupted an infrared beam within the lickport (custom-built at the UCLA Electronics shop), which triggered a solenoid valve for water delivery, all of which was controlled via a DAQ board (National Instruments USB X Series Multifunction DAQ USB-6363). The mice were required to learn to associate this water reward with the presentation of the stimulus and lick during the water reward window. If an animal was not licking during the initial days of pretrials, the experimenter would pipette tiny drops of water onto the lickport every 30 trials to coax the animal to lick. Once mice had achieved 80-85% licking rate, they were advanced to the visual discrimination task. We found no significant difference in the number of pretrial sessions it took to achieve this licking threshold between WT and *Fmr1*^-/-^ mice (WT: 3.7 ± 0.4 sessions vs. *Fmr1*^-/-^: 3.2 ± 0.3 sessions; p = 0.195; unpaired, Student’s t-test).

During the go/no-go visual discrimination task (**Fig. 1A**), sinusoidal gratings drifting at 2 different directions (orthogonal orientations) were randomly presented on the screen for 3 s. The water reward was only delivered for the preferred stimulus (45° orientation), beginning 2 s after stimulus onset, but not for the non-preferred stimulus (135° orientation) (**Fig. 1A**). Mice had to learn to discriminate between the two stimuli and to lick in anticipation of the water reward for the preferred stimulus (‘go’ trial) while withholding licking for the non-preferred stimulus (‘no-go’ trial). Licking was recorded during the entire 3 s period, though only licking occurring in the reward window was recorded. Depending on the stimulus presented, the behavioral response (licking or the lack thereof) was recorded as a “Hit”, “Miss”, “Correct Rejection” (CR), or “False Alarm” (FA) (**Fig. 1A**). An incorrect response (a “Miss” during a preferred trial or an “FA” on a non-preferred trial) resulted in a time-out period (an extension of the ITI grey screen) of 6.5 s during which the animal had to wait until the next trial. On session 6 of training, if mice had not improved in performance or reached a d’ of at least 1, the punish time was either decreased to 4.5 s if there were too many misses or increased to 9.5 s if there were too many FAs. Each training session consisted of 350 trials and only the last 100 trials were used to calculate the daily performance as the d’ statistic or discriminability index:

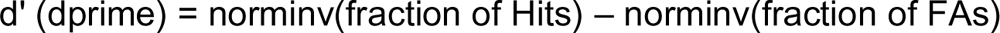

*Norminv* is a MATLAB function that returns the inverse of the normal cumulative distribution function. Custom-written routines and Psychtoolbox in MATLAB were used to deliver the visual stimuli, dispense water from the lickport, and acquire data.

### Distractor task for head-restrained mice

Once mice had maintained a d’ > 2 (the threshold we chose for expert performance) for 2 consecutive sessions (i.e., stable performance), they were advanced to the distractor task, during which auditory or visual sensory distractors were delivered in the beginning of stimulus presentation. The auditory distractor consisted of 1 beep at 5 kHz and ∼65 dB, lasting for 1.5 s and delivered from 2 speakers situated on either side of the monitor. For the visual distractor, we used LED lights (custom-made at UCLA; 580-590 nm) wrapped around the monitor (flashing 4x for 0.5 s each with a 0.25 s interstimulus interval). Distracting stimuli were delivered in only ∼50% of the trials at random and each session consisted of 200 trials. Mice performed one session of the distractor task with auditory distractors and one session of the distractor task with visual distractors on successive days; the order of the modality of distractor task was randomized. After the distractor task, another session was conducted using the standard visual discrimination task without distractors. Finally, a control session was conducted at the very end, where mice performed the task without any visual stimuli displayed on-screen to ensure that performance was dependent on stimulus presence. For two-photon calcium imaging a subset of mice were imaged at various timepoints of the training, initially before training (baseline recordings visually evoked activity without behavior) and during the distractor task.

### Human participants

Nineteen males with FXS and 20 male typically-developing healthy controls (TDC), matched on chronological age, completed the visual discrimination experiment (Table S1). 4 FXS 2 TDC females also performed the experiment. Testing was conducted at a regional academic pediatric medical center where the participants with FXS were originally recruited as part of our Center for Collaborative Research in Fragile X (U54). Approval of this study was granted through the Institutional Review Board at Cincinnati Children’s Hospital Medical Center. All participants or their legal guardians, when appropriate, provided informed written consent and/or assent prior to participating. Diagnosis of FXS was confirmed via Southern Blot and polymerase chain reaction (PCR) assays performed at Rush University in the laboratory of Dr. Elizabeth Berry-Kravis. Seven males with mosaicism (size and/or methylation) were included in all analyses unless otherwise noted. No participants had a history of non-febrile seizures or treatment with an anticonvulsant medication. Control participants were recruited through hospital-wide and community advertisements and were excluded for a history of developmental or learning disorders or significant psychiatric disorder (e.g., schizophrenia) in themselves or first degree-relatives, or for a family history of ASD in first- or second-degree relatives based on a brief screening interview. All study procedures were approved by the local Institutional Review Board.

### Visual discrimination and distractor task for human participants

Human FXS and control participants completed a visual discrimination task, followed by a distractor task that was analogous to that used with mice with relatively minor modifications. Due to the additional cognitive demands of a go/no-go paradigm, including inhibitory control, which is known to be impaired in FXS (Hooper et al., 2008), we designed a forced two-choice visual discrimination task, so that all FXS participants could learn and perform the task. Although it is possible that participants with FXS could have learned the go/no-go task with subsequent training sessions, just as the mice required consecutive sessions to learn, due to time constraints and significant burden on the patient population, this limited our ability to do so. Visual gratings were displayed via Psychtoolbox using MATLAB 2016a on a 23-inch Tobii TX300 monitor and made responses on designated keys on the keyboard. During the task, when the visual grating appeared to move from right side to left side, subjects were instructed to press the corresponding left-sided key (‘Z’), and when the visual grating appeared to move from left to right, subjects were instructed to press the corresponding right-sided key (‘M’). If participants correctly responded to the direction of the stimulus, they received positive visual feedback (e.g., image of popular video game cartoon character). If participants incorrectly responded to the direction of the stimulus, they received negative visual feedback (e.g., a large red ‘X’). If no response was received, no feedback was given. Visual gratings appeared on screen for up to 2 s or until participant response, at which point immediate feedback was presented for 1 s. Though the stimulus disappeared at 2 s, participants had until 3 s post-stimulus onset to respond and receive valid feedback. There was an intertrial interval of 2 s. All participants completed the first-order discrimination task, immediately followed by the distractor task in which auditory distractors were presented simultaneously with the visual stimuli for 50% of the trials at random.

Prior to administration of the initial task, participants received verbal instructions and then verified initial task comprehension by verbally and/or nonverbally demonstrating their expected behavioral response (i.e., pointing to left). Next, participants completed at least one block of 15 trials, in which vertical lines moved from left to right on the screen (or right to left), and participants were instructed to press the corresponding key based on the direction the lines moved. All participants included in the sample met practice criterion. Depending on the stimulus presented, the subject’s behavioral response was characterized as “Right (similar to Hit)”, “NR (no response)” or “Wrong (similar to FA)”. Since this was a forced two-choice visual discrimination task, a modified dprime, or d’ (discriminability index), was calculated as follows:

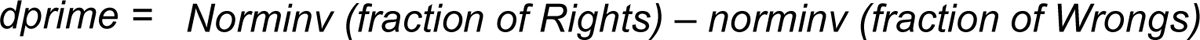

### Additional measures for human participants

All participants completed the abbreviated Stanford-Binet-Fifth Edition (SB5) to estimate general intellectual functioning. Based on previous studies(Sansone et al., 2014) we converted standard scores to Deviation IQ scores in order to reduce floor effects present for individuals with severe cognitive impairments and to better evaluate inter-individual variability.

In addition, we collected caregiver-report measures of behavior and psychiatric symptoms, including hyperactivity (Aberrant Behavior Checklist (Aman et al., 1985) and Anxiety, Depression, and Mood Scale (Esbensen et al., 2003)) as well as sensory sensitivity (Brown and Dunn, 2002) in order to relate to task performance. Participants with FXS also completed a computerized Kiddie Test of Attentional Performance (KiTAP) which has been validated for use in this population (Knox et al., 2012). KiTAP examines executive functioning through multiple subtests, including Alertness (processing speed), Distractibility (attention), Go/NoGo (response inhibition), and Flexibility (cognitive flexibility). Number of correct trials on tasks were examined in relation to correct trials on visual discrimination task with distractors. Eleven participants did not complete KiTAP due to behavior issues and/or time constraints.

Last, whole blood samples were obtained via venipuncture from participants with FXS, and analyzed using our validated Luminex-based immunoassay to determine FMRP levels (Boggs et al., 2022). Briefly, blood samples were spotted onto Whatman Bloodstain cards, and then dried blood spots were hole-punched from the cards and proteins were eluted. The eluate was analyzed in triplicate against a 9-point standard curve generated from a recombinant protein to determine participants’ FMRP concentration. One participant with FXS did not consent to blood draw.

### Support Vector Machine (SVM)

To analyze the licking data, we used the SVM available in the MATLAB Machine Learning and Deep Learning toolbox via the function *fitcsvm*. SVMs are supervised learning models with learning algorithms that can be used to classify data in a high dimensional space; it uses a subset of training points to develop a binary classifier that can use features of the data to make predictions. We used a radial basis function as the kernel. 80% of our data was applied to training the machine and 20% applied to testing it.

We performed two types of analysis using the SVM. We first binned licking into 0.1 s bins. We then used the bins prior to the water reward (0 s to 1.9 s) as the feature space of the SVM and then performed a bootstrap of 1,000 iterations in which licking was the predictor of stimulus type per mouse per session day. This generated a distribution of accuracy percentages which were then averaged. Additionally, to determine the most predictive features, we performed a bootstrapped SVM per feature of 10,000 iterations to determine the most predicative features per mouse per session day (e.g., the predictability of licking at the .1 s bin in the naive day). This again provided a distribution of accuracy percentages which were then averaged and plotted as a function of time. For each SVM set, we performed controls in which stimuli were randomly shuffled.

### Viral Constructs

Both pGP-AAV-syn-jGCaMP7f-WPRE and pGP-AAV-syn-FLEX-jGCaMP7f-WPRE were purchased from Addgene (#104488-AAV1 & #104492-AAV1) and diluted to a working titer of 1×10^13^ (to enable a longer period of optimal expression) with 1% filtered Fast Green FCF dye (Fisher Scientific). We injected (see below) a cocktail of these viruses to improve the efficacy of viral expression in pyramidal and VIP cells.

### Cranial window surgery

Craniotomies were performed at 6-8 weeks as previously described(Mostany and Portera-Cailliau, 2008; Holtmaat et al., 2009). Briefly, mice were anesthetized with isoflurane (5%, 1.5-2% maintenance), head-fixed to a stereotaxic frame and, under sterile conditions, a 4.5 mm diameter craniotomy was drilled over the V1 and covered with a 5 mm glass coverslip using cyanoacrylate glue and dental cement. Before placing the coverslip, we injected ∼60-100 nl of a cocktail of pGP-AAV-syn-jGCaMP7f-WPRE and pGP-AAV-syn-FLEX-jGCaMP7f-WPRE using a programmable nanoliter Injector (Drummond Scientific Nanoject III). A U-shaped titanium bar was attached to the skull with dental cement to head-restrain the animal during behavior and calcium imaging. For a subset of mice that underwent behavioral testing but no calcium imaging, only a head bar attachment surgery was performed (no craniotomy). All mice were administered dexamethasone (0.2 mg/Kg) i.p. or s.c. on the day of surgery to prevent swelling of the brain; a subset of mice was also administered carprofen (5 mg/Kg) as an analgesic and anti-inflammatory). After surgery, mice were placed on a heated pad for post-operative recovery until effects of the anesthesia wore off. Post-operative checks were done every 24 h for the following 2 d to ensure a healthy, full recovery.

### In vivo two-photon calcium imaging

Two-photon calcium imaging was performed on a Scientifica two-photon microscope equipped with a Chameleon Ultra II Ti:sapphire laser (Coherent) tuned to 920-940 nm, resonant scanning mirrors (Cambridge Technologies), a 20X objective (1.05 NA, Olympus), multialkali photomultiplier tubes (R3896, Hamamatsu) and ScanImage software (Pologruto et al., 2003). Prior to calcium imaging, head-restrained mice were habituated to a sound-proof chamber and allowed to run freely on a polystyrene ball and acclimated to the rig as described above for the visual task. To record visually evoked activity, we presented visual stimuli consisting of full-field sinusoidal drifting gratings (16 random repeats of 8 orientations) presented for 3 s each and separated by a 3 s-long grey screen interstimulus interval. Both spontaneous and visually evoked responses of L2/3 pyramidal cells and VIP cells from V1 were recorded at 15 Hz in 2-4 fields of view. In Fig. 3, each FOV consisted of a median of 64 pyramidal cells (range was 62-91 for WT and 30-108 cells for *Fmr1*^-/-^ mice). In Fig. 4 and Supplementary Fig. 7, each FOV consisted of a median of 4 VIP cells (range was 1-10 cells for WT and 2-10 cells for *Fmr1*^-/-^ mice). In Fig. 5 and Supplementary Figs. 8-9, each FOV consisted of a median of 6 VIP cells (range was 4-12 cells for WT and 4-13 cells for *Fmr1*^-/-^ mice). In each animal, imaging was performed at a depth of 150-200 μm, and data was averaged from movies collected across all FOVs.

### Data analysis for calcium imaging

All calcium imaging data was initially processed using *Suite2p* or *EZcalcium* software and algorithms (Pachitariu M et al., 2017; Cantu et al., 2020) for image registration, ROI detection, cell identification, and signal extraction with neuropil correction. This was done separately for pyramidal cells and VIP cells. Once *Suite2p* had performed a rigid and non-rigid registration and then detected regions-of-interest (ROIs) using a classifier, we then selected cells after visual inspection of the shape of the ROI and its fluorescence trace for quality control purposes. Next, the extracted fluorescence signal intensities for each ROI (*F*) were processed with custom-written MATLAB routines, which included modifications of our previously described code (Goel et al., 2018). A “modified Z-score” *Z_f_* (t) vector representing the activity levels of each neuron was calculated as:

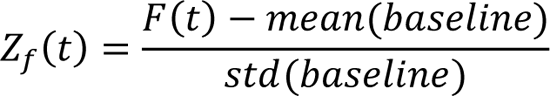

where the baseline period is the 10 s “quietest” period with the lowest variation (standard deviation) in Δ*F*/*F* (He et al., 2017) All subsequent analyses were performed using the *Z_f_* (t) vectors.

Neuropil subtraction was performed by removing the local fluorescence signal surrounding each ROI (La Fata et al., 2014). Peaks of activity were then detected in the Z-scores using the *PeakFinder* MATLAB script. These peaks were used to calculate the mean Z-score fluorescence (an estimate of amplitude of the fluorescence signal) and the frequency of events. To remove any bias resulting from peak detection, especially in VIP cells, we also calculated the frequency of events based on the magnitude of the fluorescence signal (area under the curve; AUC). For this analysis, we calculated the AUC for each fluorescence trace and divided that by the number of frames during which a stimulus occurred (i.e., 45 frames at 15 Hz for for 3 s). This was then multiplied by the frame rate to get a Z-score of fluorescence (mean activity) per second (**Fig. 4-5; Supplementary Fig. 8-9**).

To quantify visually evoked activity, we averaged the responses of neurons during the 3 s of visual stimulation and the 3 s of grey screen before the next stimulus. To quantify spontaneous activity, we conducted separate recordings during which the animals were presented a static grey screen. To determine whether an individual cell showed was responsive to visual stimuli (**Fig. 3-4**) we used a probabilistic bootstrapping method as described previously (He et al., 2017; Goel et al., 2018). First, we calculated the correlation between the stimulus time-course and the Z_F vector, followed by correlation calculations between the stimulus time-course and 1,000 scrambles of all calcium activity epochs in Z_F (epoch = consecutive frames wherein Z_F ≥ 3). The 1,000 comparisons generated a distribution of correlations (R values), within which the correlation of the unscrambled data and the stimulus fell at any given percentile. If the calculated percentile for a cell was less than 1%, then we considered that cell as being stimulus selective.

The modulation index was calculated to compare changes in activity (**Fig. 4-5; Supplementary Fig. 8-9**). In Fig. 4 we compared visually evoked activity of VIP cells to their spontaneous activity and measured the change in activity (i.e., the difference between gray screen and drifting gratings). In Fig. 5, we compared VIP activity during correct responses (hits and CRs) to activity during the majority of incorrect responses (FAs and misses). We divided mean visually evoked activity during incorrect responses by mean visually evoked activity during correct responses to get the modulation index.

### ROC analysis

The discrimination performance of pyramidal neurons was quantified using a receiver operating characteristic (ROC) analysis (O’Connor et al., 2010). Neuronal output was classified based on the similarity of response on each trial to the mean Post Stimulus Time Histograms (PSTHs) for the Hit and CR trials. Mean PSTHs were computed separately for each hit trial and CR trial. For this calculation, the current trial was not included. Similar to O’Connor et al. (2010), each trials was then assigned a “decision variable” score (DV), which was equal to the dot product similarity to the mean PSTH for Hit trials minus the dot product similarity to the mean PSTH for CR trails. Thus, DV was calculated using the following equations for Hit and CR trials respectively:

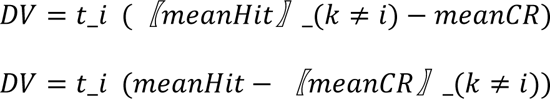

Where t_i is the PSTH_ for the current (i^th^ trial), meanHit and mean CR are the mean hit and CR PSTHs. In cases where the decision variable was large it implied a higher similarity to the mean hit PSTH compared to the mean correct rejection PSTH. IF the DV > a criterion value, the a trial was classified at “Hit”, otherwise it was a “correct rejection” trial. To determine the proportion of correctly identified trials an ROC curve was constructed(Green and Swets, 1966). The area under the ROC curve was calculated using the MATLAB function *trapz*.

### Statistical analyses

In all figures, significance levels are represented with the following convention: * for p<0.05; ** for p<0.01, *** for p<0.001. The standard error of the mean (s.e.m) is plotted using error bars unless otherwise noted. Graphs show either individual data points from each animal/human subject, or group means (average over different mice or human subjects) superimposed on individual data points. All statistical details are described in the figure legends and tests were selected based on the distribution of the data points. For parametric, two group analyses, we used a Student’s t test (paired or unpaired), and for multiple group analyses we used a one-way or two-way ANOVA. For non-parametric tests, we used the Mann-Whitney Test, Welch’s test, Friedman (repeated measures) test, and mixed-effects analysis of variance (for datasets with missing values). Multiple comparisons were corrected and correlations were conducted using Pearson’s r. For distributions, a two-sample Kolmogorov-Smirnov test was used. All statistical analyses were conducted using GraphPad Prism software or MATLAB.

### Sample Size

We determined sample sizes prior to experiments based on our past experience and the published literature, and guided by ethical principles in the use of animals (i.e., trying to minimize the number of animals used). Based on our prior studies, we estimated that for behavioral studies a minimum of 15 animals per group would be needed and, for calcium imaging, 5-6 mice would be needed. For results shown in Fig. 1, we used sample sizes of ≥ 20 mice per genotype, while in Fig. 3, we analyzed pyramidal cell data for a subset of mice (n ≥ 4 for each group of mice). We used sample sizes of > 10 for each group in Fig.4. n ≥ 5 mice per group in Fig. 5. Subsequent statistics were performed using the number of mice or subjects as the sample size or the number of cells. We chose our sample sizes for feasibility and ethical purposes in our use of animals (i.e., trying to minimize the number of animals used).

Subsequent statistics were performed using the number of mice or subjects as the sample size or the number of cells. Given the low density of VIP cells some of the analyses were done using cells as sample size, similar to other recent studies (Ren et al., 2022). However, the overall effects of reduced modulation of VIP cells on behavior was significant when mice were used as sample size (Fig. 5).

### Randomization

We ensured that during each behavior training cycle both WT and *Fmr1*^-/-^ mice were included to exclude any biases introduced by experimenters or the training rig. In addition, on a particular testing day, Fragile X participants were randomized with control subjects.

### Blinding

Experimenters were blinded to the genotype while training mice on the task. Analysis was done while being blind to the genotype.

### Neural data exclusion

We included only neurons that elicited at least one calcium transient during the duration of the recording; A small fraction of neurons was excluded because they were deemed inactive on the basis of calcium imaging data (Percentage of pyramidal neurons excluded: WT= 0.1%; *Fmr1^-/-^*= 0.1%; VIP neurons excluded: WT= 0%, *Fmr1*^-/-^ = 0.02%).

### Exclusion of mice or human participants

Two mice were excluded from the dataset because they developed health conditions or died. And two additional mice were excluded from the dataset because the mice lost more than 25% of their original body weight, which could lead to less grooming, less social interaction with cage mates, lethargy, seizures, and other health conditions that might conflict with behavior.

We excluded 5 ASD subjects from the data and one FXS participant who showed significant anomalies during the MRI session.

### Reporting summary

Further information on experimental design is available in the Life Sciences Reporting Summary linked to this article.

## ACKNOWLEDGEMENTS

The authors thank Nazim Kourdougli, Trishala Chari for help with behavioral studies. **Funding:** this work was supported by the following grants: R01NS117597 (NIH-NINDS), R01HD054453 (NIH-NICHD), Department of Defense (DOD, 13196175) awarded to C.P-C and Brain and Behavior Young Investigator Award to A.G. We also acknowledge the individuals with FXS, their families, and the controls who participated in this study. We also want to thank the clinical research coordinators for their assistance with data collection and data entry as well as the basic laboratory research assistants for their work on the FMRP assay. We also appreciate the support from Elizabeth Berry-Kravis at Rush University for testing for *FMR1* gene CGG expansion and gene methylation.

## Author contributions

A.G. and C.P-C. conceived the project and designed the experiments with help from L.d.S., L.M.S., E.P. and C.A.E. for the human studies. A.G. developed the behavioral paradigm for mice and humans. A.G., N.R, and S.P wrote the MATLAB code for analysis. A.G, N.R., L.M.S., L.d.S., J.R. and G.C. conducted the experiments. A.G., N.R. and S.P. analyzed the data. A.G., N. R., L.M.S., E.P., C.A.E. and C.P-C. interpreted the data and wrote the paper with input from other authors.

## FIGURE LEGENDS

**Supplementary Figure 1:** *Fmr1*^-/-^ mice obtained a significantly lower percentage of CR responses (27.5 ± 2.2 % for WT mice vs. 14.6 ± 2.3 % for *Fmr1*^-/-^ mice; Mann-Whitney test, p = 0.0003) and significantly higher percentage of FA responses (22.5 ± 2.2 % for WT mice vs. 35.3 ± 2.4 % for *Fmr1*^-/-^ mice; Mann-Whitney test, p *=* 0.0005) on session 4 of the visual task. There was no significant difference between genotypes in percentage of hit responses (41.6 ± 2.0 % for WT mice vs. 45.4 ± 1.3 % for *Fmr1*^-/-^ mice; Mann-Whitney test, p *=* 0.103) or percentage of miss responses (8.4 ± 2.0 % for WT mice vs. 4.7 ± 1.3 % for *Fmr1*^-/-^ mice; Mann-Whitney test, p = 0.076). Whiskers show min and max; n values are for mice, indicated on each plot. *p < 0.05; **p < 0.01; ***p < 0.001; ****p < 0.0001. (corresponds to data in Fig. 1)

**Supplementary Figure 2: A)** Illustration and timeline of behavior paradigm for visual distractor visual discrimination task. Visual distractors comprised of a string of string of LED lights flashed 4x for 0.2 s each and were presented on 50% of trials with preferred and non-preferred visual stimuli. **B)** There was no significant difference between performance (measured by the discriminability index) of WT and *Fmr1*^-/-^ mice once they reached the expert performance threshold of d’=2 on the standard visual task (2.7 ± 0.1 for WT mice vs. 3.1 ± 0.2 for *Fmr1*^-/-^ mice; Mann-Whitney test, p = 0.125, n = 19 WT mice and 19 *Fmr1*^-/-^ mice). On distractor sessions, during trials without distractors, there was no significant difference between performance of WT and *Fmr1*^-/-^ mice (2.2 ± 0.2 for WT mice vs. 1.8 ± 0.2 for *Fmr1*^-/-^ mice; Mann-Whitney test, p *=* 0.163). During trials with visual distractors, *Fmr1*^-/-^ mice exhibited worse performance than WT mice (2.1 ± 0.2 for WT mice vs. 1.4 ± 0.2 for *Fmr1*^-/-^ mice; Mann-Whitney test, p *=* 0.0097). A final session of the visual task was conducted following the distractor task; there was no significant difference between performance of WT and *Fmr1*^-/-^ mice (2.5 ± 0.1 for WT mice vs. 2.4 ± 0.2 for *Fmr1*^-/-^ mice; Mann-Whitney test, p *=* 0.515). There was no significant difference between WT performance on trials without distractors versus trials with distractors on the auditory distractor session (1.8 ± 0.2 for no light trials vs. 1.4 ± 0.2 for yes light trials; Wilcoxon matched-pairs signed rank test, p *=* 0.829). However, *Fmr1*^-/-^ mice exhibited worse performance on trials with distractors than on trials without distractors (2 ± 0.2 for no light trials vs. 1.7 ± 0.2 for yes light trials; Wilcoxon matched-pairs signed rank test, p = 0.032). **C)** *Fmr1*^-/-^ mice were more sensitive to distracting lights than WT mice. Performance (d’) was tracked throughout the last session of the visual discrimination task when learning has occurred and compared to performance throughout distractor trials on the distractor session; trials were grouped into bins of 10 trials. *Fmr1*^-/-^ performance took longer to recover to “expert” level and continued to be significantly worse than prior levels of performance throughout the duration of the task (two-way mixed ANOVA for WT basic vs. distractor; time: *F*_8,257_ = 1.4, p = 0.07; task type: *F*_1,36_ = 10, p = 0.003; two-way mixed ANOVA for *Fmr1*^-/-^ basic vs. distractor; time: *F*_7,243_ = 3.4, p = 0.002; task type: *F*_1,39_ = 18.1, p = 1.3E-4; two-way mixed ANOVA for WT learned vs. *Fmr1*^-/-^ learned; time: *F*_8,360_ = 2.7, p = 0.006; genotype: *F*_1,44_ = 0.3, p = 0.594; time x genotype: *F*_12,526_ = 1.2, p = 0.298; two-way mixed ANOVA for WT distractor vs. *Fmr1*^-/-^ distractor; time: *F*_6,164_ = 3.6, p = 0.002; genotype: *F*_1,31_ = 2.4, p = 0.013). In panels **B-C**, horizontal bars indicate mean and error bars indicate s.e.m. n values are for mice, indicated on each plot. *p < 0.05; **p < 0.01; ***p < 0.001; ****p < 0.0001. (corresponds to data in Fig. 1)

**Supplementary Figure 3: A)** *Fmr1*^-/-^ correlation between the number of sessions to reach a d’>2 on the standard visual discrimination task and the number of bins of distractor trials on the auditory distractor task with low performance (d’<2) (Pearson’s r, r = 0.4876, p = 0.025; n = 21 *Fmr1*^-/-^ mice). **B)** Correlation between d’ on distractor trials and d’ on no-distractor trials on the auditory distractor task (Pearson’s r, r = 0.8173, p = 6.1E-6; n = 21 *Fmr1*^-/-^ mice). n values are for mice, indicated on each plot. *p < 0.05; **p < 0.01; ***p < 0.001; ****p < 0.0001. (corresponds to data in Fig. 1)

**Supplementary Figure 4: A)** Performance throughout the auditory distractor task - the number of bins of trials before mice achieved expert performance (d’>2) and before they got a d’>2 twice consecutively was recorded. *Fmr1*^-/-^ mice took longer before reaching a d’>2 on a single bin (1.8 ± 0.2 for WT mice vs. 3.2 ± 0.4 for *Fmr1*^-/-^ mice; Mann-Whitney test; p = 0.003; n = 21 WT mice and 21 *Fmr1*^-/-^ mice) and longer before getting a d’>2 twice in a row than WT mice took (2.3 ± 0.4 for WT mice vs. 4.4 ± 0.7 for *Fmr1*^-/-^ mice; Mann-Whitney test; p = 0.002; n = 21 WT mice and 21 *Fmr1*^-/-^ mice). **B)** Performance throughout the visual distractor task. *Fmr1*^-/-^ mice took longer before reaching a d’>2 on a single bin (2 ± 0.4 for WT mice vs. 4.3 ± 0.7 for *Fmr1*^-/-^ mice; Mann-Whitney test; p = 0.003; n = 17 WT mice and 19 *Fmr1*^-/-^ mice) and longer before getting a d’>2 twice in a row than WT mice took (2.9 ± 0.6 for WT mice vs. 5.4 ± 0.8 for *Fmr1*^-/-^ mice; Mann-Whitney test; p = 0.016; n = 17 WT mice and 19 *Fmr1*^-/-^ mice). **C)** Genotype differences in performance (d’) are shown for each session type. There was no significant difference between WT mice and *Fmr1*^-/-^ mice on the naïve session of the visual task nor the learned session of the visual task. However, *Fmr1*^-/-^ mice exhibited worse performance on the distractor session than WT controls (Bonferroni’s multiple comparisons test for genotype effect; naïve: p = 0.083; learned: p = 0.629; distractor: p = 0.003). **D)** During the first half of the auditory distractor task, there was no difference in proportion of hit responses (0.9 ± 0.09 for WT mice vs. 0.86 ± 0.15 for *Fmr1*^-/-^ mice; Mann-Whitney test; p = 0.126), CR responses (0.65 ± 0.23 for WT mice vs. 0.54 ± 0.23 for *Fmr1*^-/-^ mice; Mann-Whitney test; p = 0.078), and miss responses (0.1 ± 0.09 for WT mice vs. 0.14 ± 0.15 for *Fmr1*^-/-^ mice; Mann-Whitney test; p = 0.126). There was a trend towards a higher percentage of FA responses (0.35 ± 0.23 for WT mice vs. 0.46 ± 0.23 for *Fmr1*^-/-^ mice; Mann-Whitney test; p = 0.078). In panels **A-D**, horizontal bars indicate mean and error bars indicate s.e.m; n values are for mice, indicated on each figure. *p < 0.05; **p < 0.01; ***p < 0.001; ****p < 0.0001. (corresponds to data in Fig. 1)

**Supplementary Figure 5: A)** Raster plots showing sample mice (n = 3 WT mice & 3 *Fmr1*^-/-^ mice) licking on trials of the auditory distractor task when distractors were present. Green is preferred and red is non-preferred. WT mice licked persistently right before and during the water window (2-3 s) on preferred trials and barely licked on non-preferred trials. *Fmr1*^-/-^ mice licked compulsively throughout most of the trial period on preferred trials and licked continuously on several of the earlier non-preferred trials in anticipation of a non-existent water reward. **B)** Graphs showing the probability of a mouse licking as a function of time during the distractor trial period. On average, probability of licking for WT mice ramped up early on preferred trials and remained relatively stable on non-preferred trials. However, for *Fmr1*^-/-^ mice it was a more gradual ramping up on preferred trials and probability also ramped up on non-preferred trials in anticipation of the water reward before going back down. Differences in licking probability between different stimulus types (preferred and non-preferred) were smaller for *Fmr1*^-/-^ mice and increased later on (two-way ANOVA; time: *F*_5,228_ = 3, p = 0.011; stim type: *F*_1,228_ = 3.9, p = 0.048; time x stimulus type: *F*_5,228_ = 2, p = 0.077; multiple paired t tests; 0-0.5 s: p = 0.013; 0.5-1 s: p = 0.942; 1-1.5 s: p = 0.043; 1.5-2 s: p = 0.013; 2-2.5 s: p = 3.6E-5; 2.5-3 s: p = 6.7E-6; n = 18 *Fmr1*^-/-^ mice) compared to WT mice (two-way ANOVA; time: *F*_5,204_ = 20.5, p < 0.0001; stim type: *F*_1,204_ = 27.5, p = 4.0E-7; time x stimulus type: *F*_5,204_ = 6, p = 3.4E-5; multiple paired t tests; 0-0.5 s: p = 0.002; 0.5-1 s: p = 0.36; 1-1.5 s: p = 0.007; 1.5-2 s: p = 3.4E-5; 2-2.5 s: p = 3.2E-6; 2.5-3 s: p = 0.004; n = 20 WT mice). *p < 0.05; **p < 0.01; ***p < 0.001; ****p < 0.0001. (corresponds to data in Fig. 1)

**Supplementary Figure 6: A)** Sex differences between performance on the visual discrimination task and distractor task in both WT and *Fmr1*^-/-^ mice were tracked. For mice performing the auditory distractor task, a mixed-effects analysis revealed a significant effect of session type and interaction effect of session type x genotype (three-way mixed ANOVA; session type: *F*_3,107_ = 22.6, p = 2.0E-11; genotype: *F*_1,38_ = 0.6, p = 0.457; sex: *F*_1,38_ = 0.1, p = 0.761; session type x genotype interaction: *F*_3,107_ = 5.6, p = 0.001; session type x sex interaction: *F*_3,107_ = 1.3, p = 0.279; genotype x sex interaction: *F*_1,38_ = 1.2, p = 0.283; session type x genotype x sex interaction: *F*_3,107_ = 1.7, p = 0.171). There was no significant difference between performance of WT female and male mice once they reached the expert performance threshold of d’=2 on the visual task (2.8 ± 0.2 for WT females and 3.8 ± 0.1 for WT males; Mann-Whitney test, p = 0.927; n = 7 WT females and 14 WT males). In addition, there was no significant difference between performance of *Fmr1*^-/-^ female and male mice once they reached the expert performance threshold of d’=2 on the visual task (3.1 ± 0.2 for *Fmr1*^-/-^ females and 3.4 ± 0.3 for *Fmr1*^-/-^ males; Mann-Whitney test, p = 0.784; n = 7 *Fmr1*^-/-^ females and 14 *Fmr1*^-/-^ males).On auditory distractor sessions, during trials without distractors, there was no significant difference between performance of WT female and male mice (2.1 ± 0.2 for WT females and 2.5 ± 0.2 for WT males; two Mann-Whitney test, p = 0.255) and no significant difference between performance of *Fmr1*^-/-^ female and male mice (2.2 ± 0.4 for *Fmr1*^-/-^ females and 1.9 ± 0.2 for *Fmr1*^-/-^ males; Mann-Whitney test, p *=* 0.535). During trials with distractors, WT female mice performed worse than WT male mice (2 ± 0.2 for WT females and 2.6 ± 0.2 for WT males; Mann-Whitney test, p = 0.031). There was no significant difference between performance of *Fmr1*^-/-^ female and male mice (1.7 ± 0.3 for *Fmr1*^-/-^ females and 1.7 ± 0.2 for *Fmr1*^-/-^ males; Mann-Whitney test, p = 0.856). On the final session of the visual task conducted following the distractor task, there was no significant difference between performance of WT female and male mice (2.5 ± 0.2 for WT females and 2.5 ± 0.1 for WT males; Mann-Whitney test, p = 0.902) and no significant difference between performance of *Fmr1*^-/-^ female and male mice (2.8 ± 0.2 for *Fmr1*^-/-^ females and 2.2 ± 0.3 for *Fmr1*^-/-^ males; Mann-Whitney test, p = 0.147). **B)** For mice performing the visual distractor task, a mixed-effects analysis revealed a significant effect of session type, interaction effect of session type x genotype, and interaction effect of session type x genotype x sex (three-way mixed ANOVA; session type: *F*_3,98_ = 26.1, p = 1.1E-11; genotype: *F*_1,34_ = 0.7, p = 0.419; sex: *F*_1,34_ = 0.004, p = 0.948; session type x genotype interaction: *F*_3,98_ = 3.9, p = 0.011; session type x sex interaction: *F*_3,98_ = 0.9, p = 0.433; genotype x sex interaction: *F*_1,34_ = 3.2, p = 0.081; session type x genotype x sex interaction: *F*_3,98_ = 3.1, p = 0.028). There was no significant difference between performance of WT female and male mice once they reached the expert performance threshold of d’=2 on the visual task (2.8 ± 0.2 for WT females and 2.7 ± 0.1 for WT males; Mann-Whitney test, p = 0.711; n = 7 WT females and 12 WT males). In addition, there was no significant difference between performance of *Fmr1*^-/-^ female and male mice once they reached the expert performance threshold of d’=2 on the visual task (3 ± 0.1 for *Fmr1*^-/-^ females and 3.1 ± 0.3 for *Fmr1*^-/-^ males; Mann-Whitney test, p = 0.837; n = 7 *Fmr1*^-/-^ females and 12 *Fmr1*^-/-^ males). On visual distractor sessions, during trials without distractors, WT female mice performed worse than WT male mice (1.8 ± 0.3 for WT females and 2.5 ± 0.2 for WT males; Mann-Whitney test, p = 0.045). There was no significant difference between performance of *Fmr1*^-/-^ female and male mice (2.3 ± 0.3 for *Fmr1*^-/-^ females and 1.6 ± 0.2 for *Fmr1*^-/-^ males; Mann-Whitney test, p = 0.1). During trials with distractors, there was no significant difference between performance of WT female and male mice (1.7 ± 0.2 for WT females and 2.4 ± 0.2 for WT males; Mann-Whitney test, p = 0.167) and no significant difference between performance of *Fmr1*^-/-^ female and male mice (1.6 ± 0.3 for *Fmr1*^-/-^ females and 1.3 ± 0.3 for *Fmr1*^-/-^ males; Mann-Whitney test, p = 0.606). On the final session of the visual task conducted following the distractor task, there was no significant difference between performance of WT female and male mice (2.5 ± 0.2 for WT females and 2.6 ± 0.1 for WT males; Mann-Whitney test, p = 0.773) and no significant difference between performance of *Fmr1*^-/-^ female and male mice (2.7 ± 0.2 for *Fmr1*^-/-^ females and 2.2 ± 0.3 for *Fmr1*^-/-^ males; Mann-Whitney test, p = 0.181). In panels **A-B**, horizontal bars indicate mean and error bars indicate s.e.m; n values are for mice, indicated on each figure. *p < 0.05; **p < 0.01; ***p < 0.001; ****p < 0.0001. (corresponds to data in Fig. 1 and Supplementary Fig. 2)

**Supplementary Figure 7: A)** There was no significant difference between the total number of active VIP cells per field-of-view (FOV) in WT and *Fmr1*^-/-^ mice (4.2 ± 2.3 for WT mice vs. 5.3 ± 2.5 for *Fmr1*^-/-^ mice; Mann-Whitney test, p = 0.162; n = 13 WT mice and 12 *Fmr1*^-/-^ mice). In panel **A**, horizontal bars indicate mean and error bars indicate s.e.m; n values are for mice, indicated on the figure. *p < 0.05; **p < 0.01; ***p < 0.001; ****p < 0.0001. (corresponds to data in Fig. 4)

**Supplementary Figure 8: A)** During the stimulus period (0-3 s) of trials without distractors, there was reduced mean VIP activity per second during correct trials for *Fmr1*^-/-^ mice (58.5 ± 38.5 for WT vs. 54 ± 71.4 for *Fmr1*^-/-^; Mann-Whitney test, p = 0.043, Cohen’s *d* = 0.078; n = 6 WT mice, 42 VIP cells and 5 *Fmr1*^-/-^ mice, 44 VIP cells). **B)** There was even greater reduced mean VIP activity per second during error trials for *Fmr1*^-/-^ mice (40.3 ± 25.2 for WT vs. 21.3 ± 23.2 for *Fmr1*^-/-^; Mann-Whitney test, p = 6.3E-5, Cohen’s *d* = 0.784). **C)** A cumulative probability plot showing reduced VIP cell modulation by errors for *Fmr1*^-/-^ mice as measured by the modulation index (two-sample Kolmogorov-Smirnov test, p = 0.001, k = 0.4004). Bar graph inset showing VIP cells were less modulated by errors in *Fmr1*^-/-^ mice (0.8 ± 0.6 for WT vs. 0.7 ± 0.9 for *Fmr1*^-/-^; Mann-Whitney test, p = 0.0001, Cohen’s *d* = 0.214). **D)** During the post-stimulus period (3-6 s for correct trials and 3-12.5 s for error trials) of auditory distractor trials, there was no significant difference in mean VIP activity per second during correct trials (64.8 ± 38.5 for WT vs. 63.0 ± 80.5 for *Fmr1*^-/-^; Mann-Whitney test, p = 0.113). **E)** There was reduced mean VIP activity per second during error trials for *Fmr1*^-/-^ mice (109.3 ± 75.3 for WT vs. 72.3 ± 80.5 for *Fmr1*^-/-^; Mann-Whitney test, p = 0.006, Cohen’s *d* = 0.475). **F)** A cumulative probability plot showing VIP cell modulation by errors as measured by the modulation index (two-sample Kolmogorov-Smirnov test, p = 0.055, k = 0.2803). Bar graph inset showing VIP cells were less modulated by errors in *Fmr1*^-/-^ mice (1.7 ± 0.8 for WT vs. 1.3 ± 0.6 for *Fmr1*^-/-^; Mann-Whitney test, p = 0.005, Cohen’s *d* = 0.611). In panels **A-F,** horizontal bars indicate mean and error bars indicate s.e.m; n values are for mice and cells, indicated on the figure. *p < 0.05; **p < 0.01; ***p < 0.001; ****p < 0.0001. (corresponds to data in Fig. 5)

**Supplementary Figure 9: A)** During the stimulus period of distractor trials, there was reduced mean VIP activity per second during correct trials for *Fmr1*^-/-^ mice during the first 20 and middle 20 trials (multiple Mann-Whitney tests; first 20: 47.0 ± 34.8 for WT vs. 26.8 ± 38.9 for *Fmr1*^-/-^; p = 0.004, Cohen’s *d* = 0.546; middle 20: 75.9 ± 81.0 for WT vs. 45.0 ± 87.2 for *Fmr1*^-/-^; p = 0.008, Cohen’s *d* = 0.367; last 20: 47.0 ± 34.8 for WT vs. 26.8 ± 38.9 for *Fmr1*^-/-^; p = 0.233; n = 6 WT mice, 42 VIP cells and 5 *Fmr1*^-/-^ mice, 44 VIP cells). There was a significant effect of time (two-way ANOVA; time: *F*_2,168_ = 3.1, p = 0.048; genotype: *F*_1,84_ = 1.1, p = 0.297; time x genotype: *F*_2,168_ = 3, p = 0.054). **B)** During the stimulus period of distractor trials, there was even greater reduced mean VIP activity per second during error trials for *Fmr1*^-/-^ mice during the first 20 and middle 20 trials (multiple Mann-Whitney tests; first 20: 67.9 ± 72.7 for WT vs. 18.9 ± 19.9 for *Fmr1*^-/-^; p = 7.9E-5, Cohen’s *d* = 0.919; middle 20: 42.1 ± 31.9 for WT vs. 21.3 ± 28 for *Fmr1*^-/-^; p = 3.3E-4, Cohen’s *d* = 0.694; last 20: 30.0 ± 44.9 for WT vs. 21.9 ± 31.9 for *Fmr1*^-/-^; p = 0.403). There was a significant effect of time and effect of genotype (two-way mixed ANOVA; time: *F*_2,164_ = 5.7, p = 0.004; genotype: *F*_1,84_ = 15.3, p = 2.0E-4; time x genotype: *F*_2,164_ = 7.7, p = 0.054). **C)** During the post-stimulus period of distractor trials, there was reduced mean VIP activity per second during correct trials for *Fmr1*^-/-^ mice during the middle 20 trials (multiple Mann-Whitney tests; first 20: 53.2 ± 42.2 for WT vs. 46.4 ± 62.7 for *Fmr1*^-/-^; p = 0.216; middle 20: 72.4 ± 73.2 for WT vs. 56.6 ± 92.4 for *Fmr1*^-/-^; p = 0.008, Cohen’s *d* = 0.19; last 20: 41.7 ± 43.9 for WT vs. 59.8 ± 115.8 for *Fmr1*^-/-^; p = 0.183. There was no significant effect or interaction effect (two-way ANOVA; time: *F*_2,168_ = 1.6, p = 0.202; genotype: *F*_1,84_ = 1.9, p = 0.171; time x genotype: *F*_2,168_ = 1, p = 0.358). **D)** During the post-stimulus period of distractor trials, there was reduced mean VIP activity per second during error trials for *Fmr1*^-/-^ mice during the middle 20 trials (multiple Mann-Whitney tests; first 20: 154.0 ± 128.5 for WT vs. 56.8 ± 78.4 for *Fmr1*^-/-^; p = 7.9E-5, Cohen’s *d* = 0.913; middle 20: 119.0 ± 87.0 for WT vs. 62.9 ± 98.3 for *Fmr1*^-/-^; p = 3.3E-4, Cohen’s *d* = 0.605; last 20: 59.2 ± 57.6 for WT vs. 79.0 ± 116.7 for *Fmr1*^-/-^; p = 0.403. There was a significant effect of time, effect of genotype, and time x genotype interaction effect (two-way mixed ANOVA; time: *F*_2,164_ = 5.7, p = 0.004; genotype: *F*_1,84_ = 15.3, p = 2.0E-4; time x genotype: *F*_2,164_ = 7.7, p = 7.0E-4). **E)** A cumulative probability plot showing reduced VIP cell modulation by errors for *Fmr1*^-/-^ mice during the stimulus period on the first 20 distractor trials (two-sample Kolmogorov-Smirnov test, p = 3.1E-4, k = 0.438) and no significant difference on the middle 20 distractor trials (two-sample Kolmogorov-Smirnov test, p = 0.365, k = 0.193). Bar graph inset showing VIP cells were less modulated by errors in *Fmr1*^-/-^ mice on the first 20 distractor trials (1.5 ± 1.9 for WT vs. 0.7 ± 0.5 for *Fmr1*^-/-^; Mann-Whitney test, p = 4.30E-4, Cohen’s *d* = 0.569) and there was no significant difference on the middle 20 distractor trials (1.2 ± 2.3 for WT vs. 0.9 ± 1.1 for *Fmr1*^-/-^; Mann-Whitney test, p = 0.494). There was a significant effect of genotype (two-way ANOVA; time: *F*_1,84_ = 0.1, p = 0.748; genotype: *F*_1,84_ = 5.4, p = 0.022; time x genotype: *F*_1,80_ = 0.9, p = 0.33). **F)** The modulation index of VIP cells during the stimulus period of all auditory distractor trials was averaged per mouse and was trending towards a negative correlation with percent incorrect (%) on the distractor task (Pearson’s r, r = −0.4999, p = 0.059). A regression line is fitted to the data points. **G)** The modulation index of VIP cells during the stimulus period of all auditory distractor trials was trending towards a negative correlation with number of days to learn the standard visual task (Pearson’s r, r = −0.5189, p = 0.051). In panels **A-E,** horizontal bars indicate mean and error bars indicate s.e.m; n values are for mice and cells, indicated on the figure. *p < 0.05; **p < 0.01; ***p < 0.001; ****p < 0.0001. (corresponds to data in Fig. 5)

## SUPPLEMENTARY DATA

**Table S1.** Demographic Information for FXS and TDC Groups

|  | ***FXS (n=23)*** | ***TDC (n=22)*** |
| --- | --- | --- |
| *Age* | 28.7 (10.0)<br>11 - 45 | 30.5 (9.2)<br>16 - 46 |
| *Deviation IQ* | 44.4 (20.6)<sup>***</sup><br>18 - 91 | 109.6 (11.1)<br>91 - 128 |
| *Vineland ABC* | 48.6 (22.7)<br>20 - 88 | - |
| *FMRP* | 1.8 (2.4)<sup>***</sup><br>0 - 25 | 24.6 (4.8)<br>17 - 35 |
| *Sex (n, %)* | 4, 84 |  |
| *Mosaic (n, %)* | 7, 37 |  |

Mean (Standard Deviation), *Range* given unless otherwise specified
IQ – Intelligence Quotient; ABC – Adaptive Behavior Composite; FMRP – Fragile X Messenger Ribonucleoprotein; Mosaic – Size mosaicism and/or methylation mosaicism
\*\*\* p < .001

