## Supplementary Figures for "Hypersensitivity to distractors in Fragile X syndrome from loss of modulation of cortical VIP interneurons"

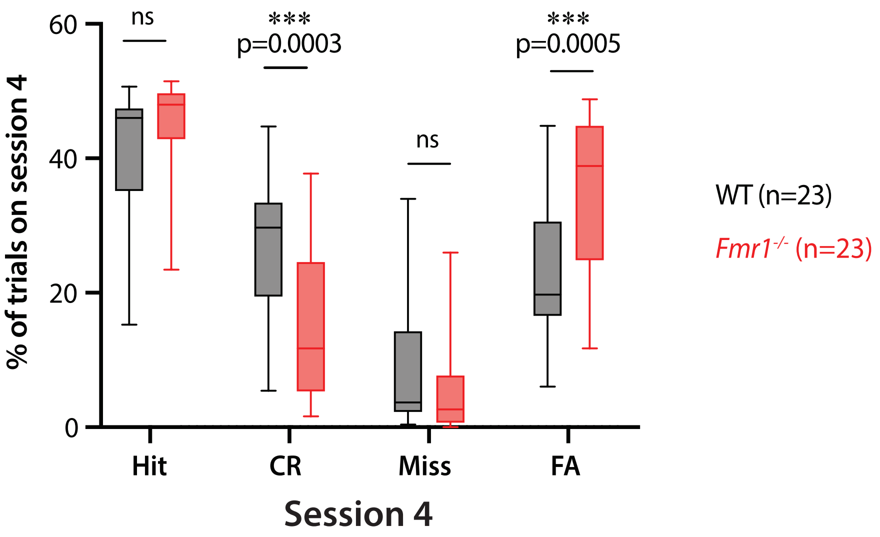


**Supplementary Figure 1:**
*Fmr1*$^{-/-}$ mice obtained a significantly lower percentage of CR responses (27.5 ± 2.2 % for WT mice vs. 14.6 ± 2.3 % for *Fmr1*$^{-/-}$ mice; Mann-Whitney test, p = 0.0003) and significantly higher percentage of FA responses (22.5 ± 2.2 % for WT mice vs. 35.3 ± 2.4 % for *Fmr1*$^{-/-}$ mice; Mann-Whitney test, p = 0.0005) on session 4 of the visual task. There was no significant difference between genotypes in percentage of hit responses (41.6 ± 2.0 % for WT mice vs. 45.4 ± 1.3 % for *Fmr1*$^{-/-}$ mice; Mann-Whitney test, p = 0.103) or percentage of miss responses (8.4 ± 2.0 % for WT mice vs. 4.7 ± 1.3 % for *Fmr1*$^{-/-}$ mice; Mann-Whitney test, p = 0.076).
Whiskers show min and max; n values are for mice, indicated on each plot. *p < 0.05; **p < 0.01; ***p < 0.001; ****p < 0.0001.
(corresponds to data in Fig. 1)

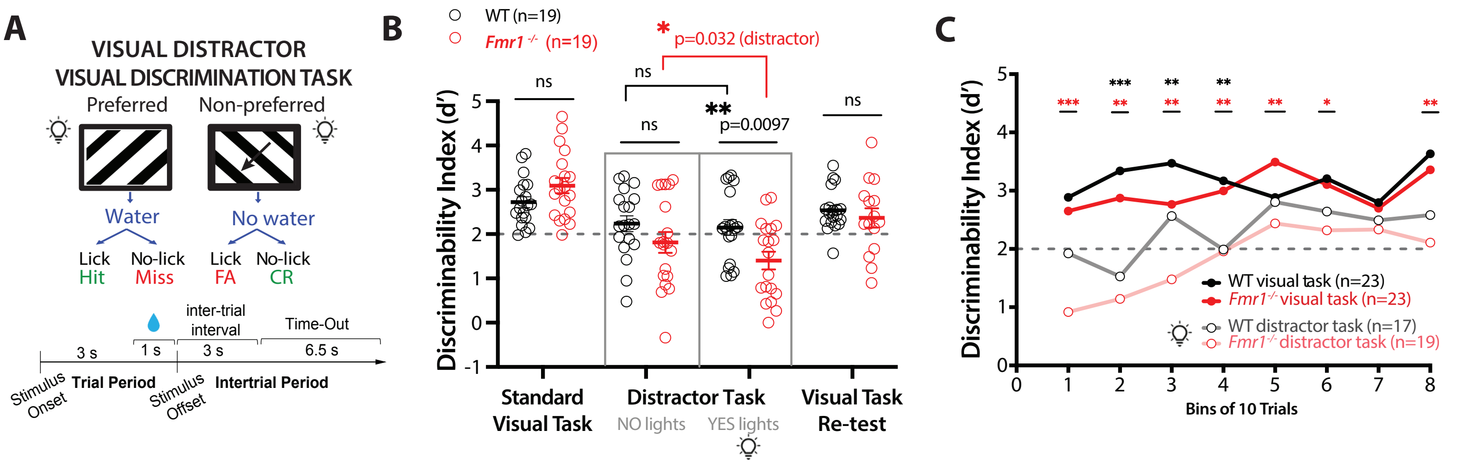


**Supplementary Figure 2:**

**A)** Illustration and timeline of behavior paradigm for visual distractor visual discrimination task. Visual distractors comprised of a string of string of LED lights flashed 4x for 0.2 s each and were presented on 50% of trials with preferred and non-preferred visual stimuli. **B)** There was no significant difference between performance (measured by the discriminability index) of WT and *Fmr1*$^{-/-}$ mice once they reached the expert performance threshold of d'=2 on the standard visual task (2.7 ± 0.1 for WT mice vs. 3.1 ± 0.2 for *Fmr1*$^{-/-}$ mice; Mann-Whitney test, p = 0.125, n = 19 WT mice and 19 Fmr1-/- mice). On distractor sessions, during trials without distractors, there was no significant difference between performance of WT and *Fmr1*$^{-/-}$ mice (2.2 ± 0.2 for WT mice vs. 1.8 ± 0.2 for *Fmr1*$^{-/-}$ mice; Mann-Whitney test, p = 0.163). During trials with visual distractors, *Fmr1*$^{-/-}$ mice exhibited worse performance than WT mice (2.1 ± 0.2 for WT mice vs. 1.4 ± 0.2 for *Fmr1*$^{-/-}$ mice; Mann-Whitney test, p = 0.0097). A final session of the visual task was conducted following the distractor task; there was no significant difference between performance of WT and *Fmr1*$^{-/-}$ mice (2.5 ± 0.1 for WT mice vs. 2.4 ± 0.2 for *Fmr1*$^{-/-}$ mice; Mann-Whitney test, p = 0.515). There was no significant difference between WT performance on trials without distractors versus trials with distractors on the auditory distractor session (1.8 ± 0.2 for no light trials vs. 1.4 ± 0.2 for yes light trials; Wilcoxon matched-pairs signed rank test, p = 0.829). However, *Fmr1*$^{-/-}$ mice exhibited worse performance on trials with distractors than on trials without distractors (2 ± 0.2 for no light trials vs. 1.7 ± 0.2 for yes light trials; Wilcoxon matched-pairs signed rank test, p = 0.032).

**C)** *Fmr1*$^{-/-}$ mice were more sensitive to distracting lights than WT mice. Performance (d') was tracked throughout the last session of the visual discrimination task when learning has occurred and compared to performance throughout distractor trials on the distractor session; trials were grouped into bins of 10 trials. *Fmr1*$^{-/-}$ performance took longer to recover to "expert" level and continued to be significantly worse than prior levels of performance throughout the duration of the task (two-way mixed ANOVA for WT basic vs. distractor; time: $F_{8,257}$ = 1.4, p = 0.07; task type: $F_{1,36}$ = 10, p = 0.003; two-way mixed ANOVA for *Fmr1*$^{-/-}$ basic vs. distractor; time: $F_{7,243}$ = 3.4, p = 0.002; task type: $F_{1,39}$ = 18.1, p = 1.3E-4; two-way mixed ANOVA for WT learned vs. *Fmr1*$^{-/-}$ learned; time: $F_{8,360}$ = 2.7, p = 0.006; genotype: $F_{1,44}$ = 0.3, p = 0.594; time x genotype: $F_{12,526}$ = 1.2, p = 0.298; two-way mixed ANOVA for WT distractor vs. *Fmr1*$^{-/-}$ distractor; time: $F_{6,164}$ = 3.6, p = 0.002; genotype: $F_{1,31}$ = 2.4, p = 0.013).

In panels B-C, horizontal bars indicate mean and error bars indicate s.e.m. n values are for mice, indicated on each plot. *p < 0.05; **p < 0.01; ***p < 0.001; ****p < 0.0001.

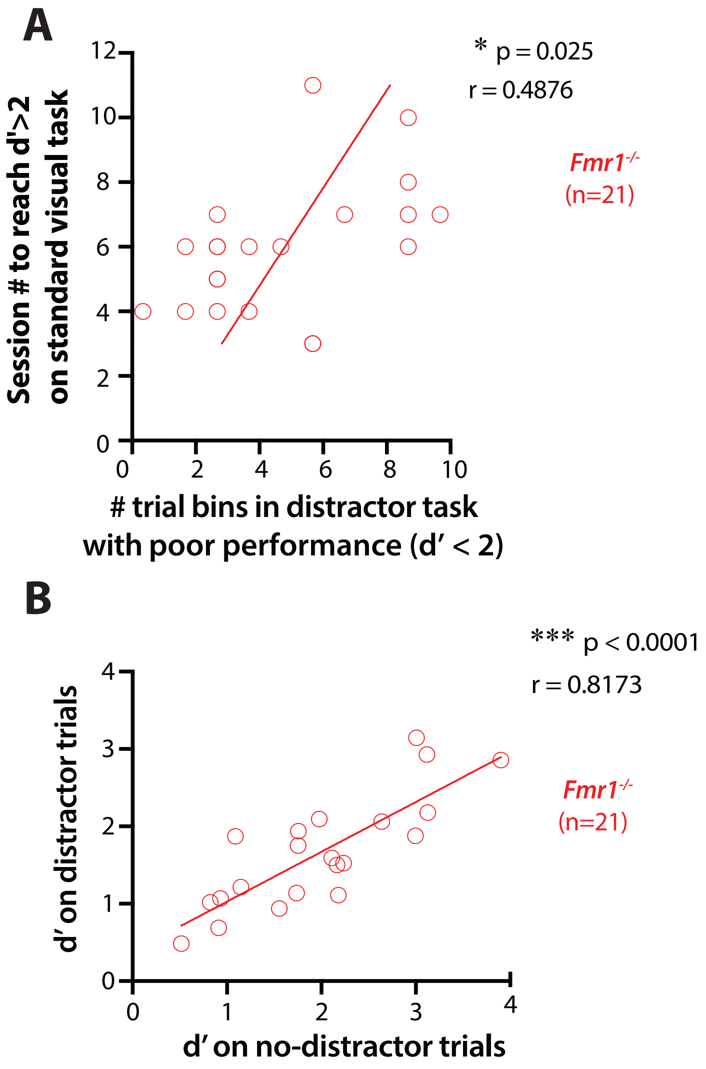


**Supplementary Figure 3:**

**A)** *Fmr1*$^{-/-}$ correlation between the number of sessions to reach a d'>2 on the standard visual discrimination task and the number of bins of distractor trials on the auditory distractor task with low performance (d'<2) (Pearson's r, r = 0.4876, p = 0.025; n = 21 *Fmr1*$^{-/-}$ mice). **B)** Correlation between d' on distractor trials and d' on no-distractor trials on the auditory distractor task (Pearson's r, r = 0.8173, p = 6.1E-6; n = 21 *Fmr1*$^{-/-}$ mice).

n values are for mice, indicated on each plot. *p < 0.05; **p < 0.01; ***p < 0.001; ****p < 0.0001.

(corresponds to data in Fig. 1)

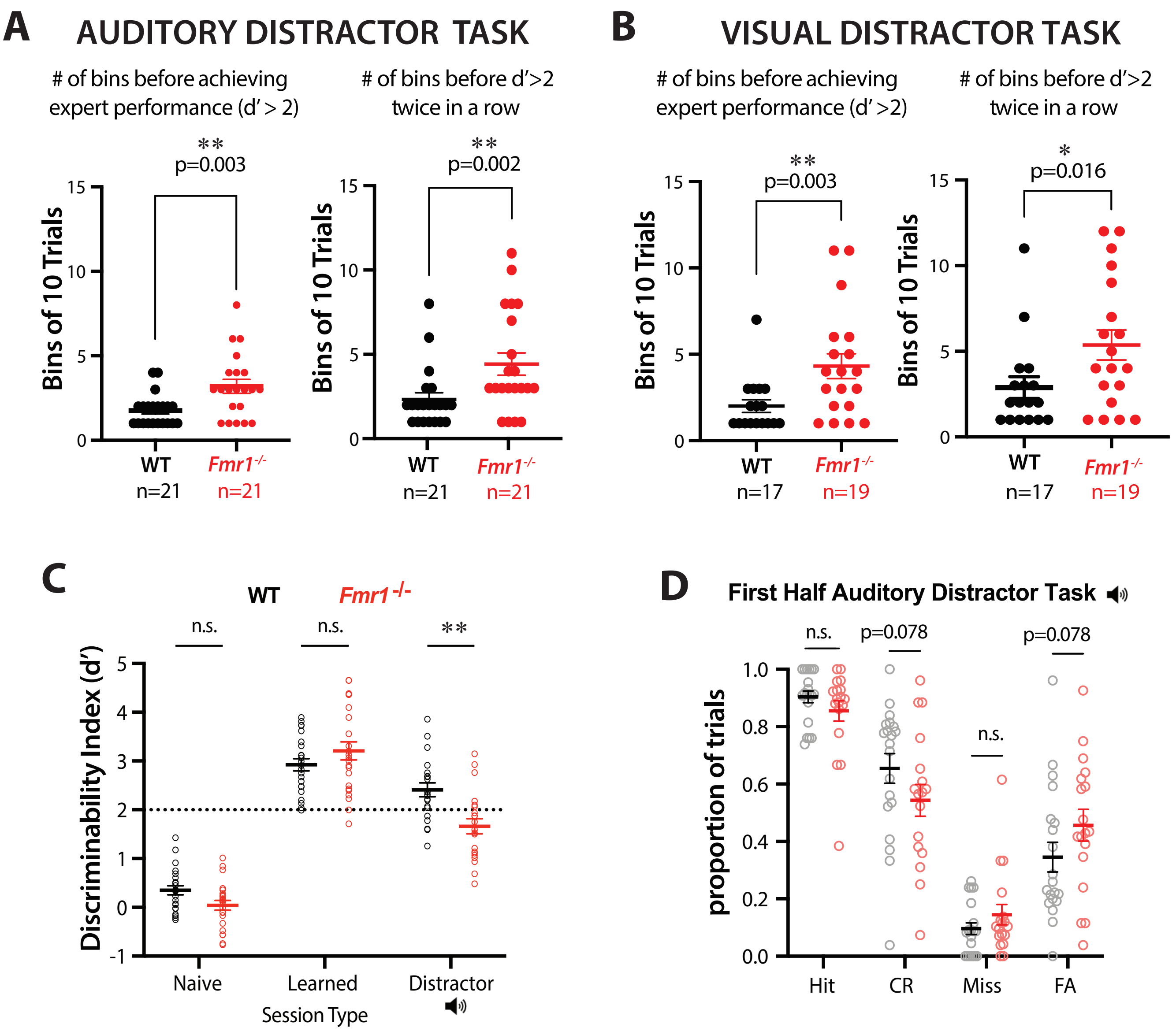


**Supplementary Figure 4:**

**A)** Performance throughout the auditory distractor task - the number of bins of trials before mice achieved expert performance (d'>2) and before they got a d'>2 twice consecutively was recorded. *Fmr1*$^{-/-}$ mice took longer before reaching a d'>2 on a single bin (1.8 ± 0.2 for WT mice vs. 3.2 ± 0.4 for *Fmr1*$^{-/-}$ mice; Mann-Whitney test; p = 0.003; n = 21 WT mice and 21 *Fmr1*$^{-/-}$ mice) and longer before getting a d'>2 twice in a row than WT mice took (2.3 ± 0.4 for WT mice vs. 4.4 ± 0.7 for *Fmr1*$^{-/-}$ mice; Mann-Whitney test; p = 0.002; n = 21 WT mice and 21 *Fmr1*$^{-/-}$ mice).

**B)** Performance throughout the visual distractor task. *Fmr1*$^{-/-}$ mice took longer before reaching a d'>2 on a single bin (2 ± 0.4 for WT mice vs. 4.3 ± 0.7 for Fmr1-/- mice; Mann-Whitney test; p = 0.003; n = 17 WT mice and 19 *Fmr1*$^{-/-}$ mice) and longer before getting a d'>2 twice in a row than WT mice took (2.9 ± 0.6 for WT mice vs. 5.4 ± 0.8 for *Fmr1*$^{-/-}$ mice; Mann-Whitney test; p = 0.016; n = 17 WT mice and 19 *Fmr1*$^{-/-}$ mice).

**Supplementary Figure 4 Legend continued:**

**C)** Genotype differences in performance (d') are shown for each session type. There was no significant difference between WT mice and *Fmr1*$^{-/-}$ mice on the naïve session of the visual task nor the learned session of the visual task. However, *Fmr1*$^{-/-}$ mice exhibited worse performance on the distractor session than WT controls (Bonferroni's multiple comparisons test for genotype effect; naïve: $p = 0.083$; learned: $p = 0.629$; distractor: $p = 0.003$).

**D)** During the first half of the auditory distractor task, there was no difference in proportion of hit responses (0.9 ± 0.09 for WT mice vs. 0.86 ± 0.15 for *Fmr1*$^{-/-}$ mice; Mann-Whitney test; $p = 0.126$), CR responses (0.65 ± 0.23 for WT mice vs. 0.54 ± 0.23 for *Fmr1*$^{-/-}$ mice; Mann-Whitney test; $p = 0.078$), and miss responses (0.1 ± 0.09 for WT mice vs. 0.14 ± 0.15 for *Fmr1*$^{-/-}$ mice; Mann-Whitney test; $p = 0.126$). There was a trend towards a higher percentage of FA responses (0.35 ± 0.23 for WT mice vs. 0.46 ± 0.23 for *Fmr1*$^{-/-}$ mice; Mann-Whitney test; $p = 0.078$).

In panels A-D, horizontal bars indicate mean and error bars indicate s.e.m; n values are for mice, indicated on each figure. *$p < 0.05$; **$p < 0.01$; ***$p < 0.001$; ****$p < 0.0001$.

(corresponds to data in Fig. 1)

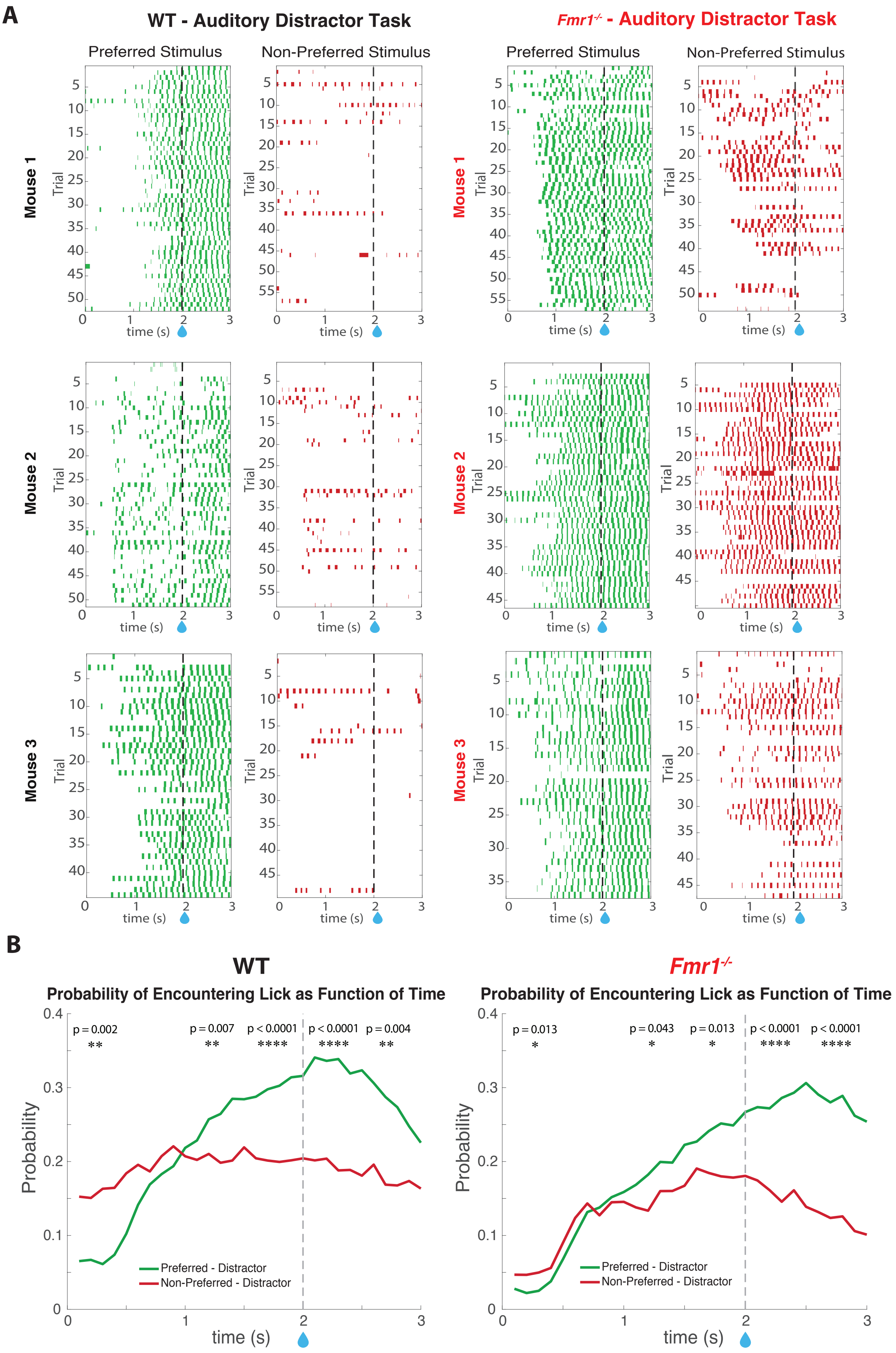


### Supplementary Figure 5

**Supplementary Figure 5 Legend:**

**A)** Raster plots showing sample mice (n = 3 WT mice & 3 *Fmr1*$^{-/-}$ mice) licking on trials of the auditory distractor task when distractors were present. Green is preferred and red is non-preferred. WT mice licked persistently right before and during the water window (2-3 s) on preferred trials and barely licked on non-preferred trials. *Fmr1*$^{-/-}$ mice licked compulsively throughout most of the trial period on preferred trials and licked continuously on several of the earlier non-preferred trials in anticipation of a non-existent water reward. **B)** Graphs showing the probability of a mouse licking as a function of time during the distractor trial period. On average, probability of licking for WT mice ramped up early on preferred trials and remained relatively stable on non-preferred trials. However, for *Fmr1*$^{-/-}$ mice it was a more gradual ramping up on preferred trials and probability also ramped up on non-preferred trials in anticipation of the water reward before going back down. Differences in licking probability between different stimulus types (preferred and non-preferred) were smaller for *Fmr1*$^{-/-}$ mice and increased later on (two-way ANOVA; time: $F_{5,228} = 3$, $p = 0.011$; stim type: $F_{1,228} = 3.9$, $p = 0.048$; time x stimulus type: $F_{5,228} = 2$, $p = 0.077$; multiple paired t tests; 0-0.5 s: $p = 0.013$; 0.5-1 s: $p = 0.942$; 1-1.5 s: $p = 0.043$; 1.5-2 s: $p = 0.013$; 2-2.5 s: $p = 3.6E-5$; 2.5-3 s: $p = 6.7E-6$; n = 18 *Fmr1*$^{-/-}$ mice) compared to WT mice (two-way ANOVA; time: $F_{5,204} = 20.5$, $p < 0.0001$; stim type: $F_{1,204} = 27.5$, $p = 4.0E-7$; time x stimulus type: $F_{5,204} = 6$, $p = 3.4E-5$; multiple paired t tests; 0-0.5 s: $p = 0.002$; 0.5-1 s: $p = 0.36$; 1-1.5 s: $p = 0.007$; 1.5-2 s: $p = 3.4E-5$; 2-2.5 s: $p = 3.2E-6$; 2.5-3 s: $p = 0.004$; n = 20 WT mice).

*p < 0.05; **p < 0.01; ***p < 0.001; ****p < 0.0001.

(corresponds to data in Fig. 1)

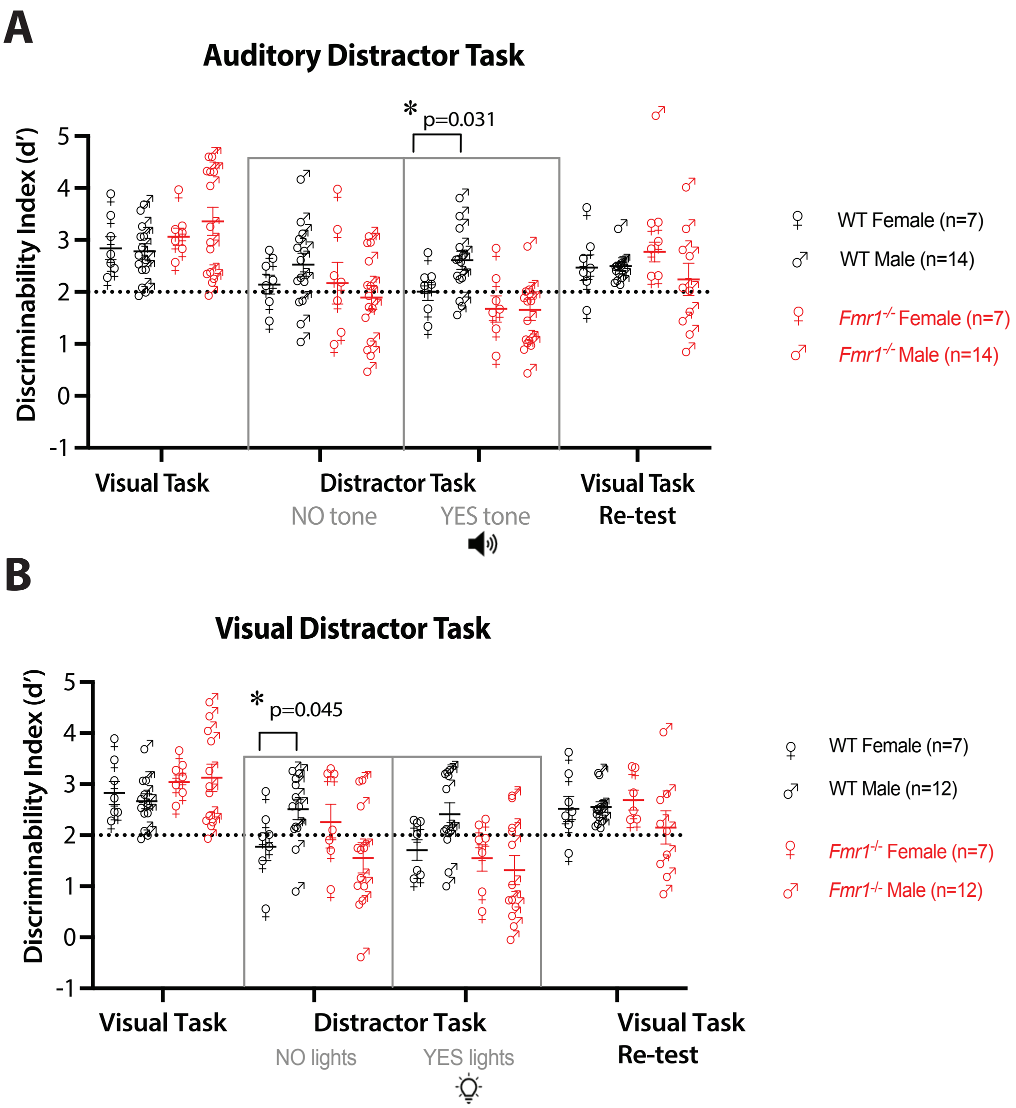


**Supplementary Figure 6**

**Supplementary Figure 6 Legend:**

**A)** Sex differences between performance on the visual discrimination task and distractor task in both WT and *Fmr1*$^{-/-}$ mice were tracked. For mice performing the auditory distractor task, a mixed-effects analysis revealed a significant effect of session type and interaction effect of session type x genotype (three-way mixed ANOVA; session type: $F_{3,107} = 22.6$, p = 2.0E-11; genotype: $F_{1,38} = 0.6$, p = 0.457; sex: F1,38 = 0.1, p = 0.761; session type x genotype interaction: $F_{3,107} = 5.6$, p = 0.001; session type x sex interaction: $F_{3,107} = 1.3$, p = 0.279; genotype x sex interaction: $F_{1,38} = 1.2$, p = 0.283; session type x genotype x sex interaction: $F_{3,107} = 1.7$, p = 0.171). There was no significant difference between performance of WT female and male mice once they reached the expert performance threshold of d'=2 on the visual task (2.8 ± 0.2 for WT females and 3.8 ± 0.1 for WT males; Mann-Whitney test, p = 0.927; n = 7 WT females and 14 WT males). In addition, there was no significant difference between performance of *Fmr1*$^{-/-}$ female and male mice once they reached the expert performance threshold of d'=2 on the visual task (3.1 ± 0.2 for *Fmr1*$^{-/-}$ females and 3.4 ± 0.3 for *Fmr1*$^{-/-}$ males; Mann-Whitney test, p = 0.784; n = 7 *Fmr1*$^{-/-}$ females and 14 *Fmr1*$^{-/-}$ males). On auditory distractor sessions, during trials without distractors, there was no significant difference between performance of WT female and male mice (2.1 ± 0.2 for WT females and 2.5 ± 0.2 for WT males; two Mann-Whitney test, p = 0.255) and no significant difference between performance of *Fmr1*$^{-/-}$ female and male mice (2.2 ± 0.4 for*Fmr1*$^{-/-}$ females and 1.9 ± 0.2 for *Fmr1*$^{-/-}$males; Mann-Whitney test, p = 0.535). During trials with distractors, WT female mice performed worse than WT male mice (2 ± 0.2 for WT females and 2.6 ± 0.2 for WT males; Mann-Whitney test, p = 0.031). There was no significant difference between performance of *Fmr1*$^{-/-}$ female and male mice (1.7 ± 0.3 for Fmr1-/- females and 1.7 ± 0.2 for *Fmr1*$^{-/-}$ males; Mann-Whitney test, p = 0.856). On the final session of the visual task conducted following the distractor task, there was no significant difference between performance of WT female and male mice (2.5 ± 0.2 for WT females and 2.5 ± 0.1 for WT males; Mann-Whitney test, p = 0.902) and no significant difference between performance of *Fmr1*$^{-/-}$ female and male mice (2.8 ± 0.2 for *Fmr1*$^{-/-}$ females and 2.2 ± 0.3 for *Fmr1*$^{-/-}$ males; Mann-Whitney test, p = 0.147).

**B)** For mice performing the visual distractor task, a mixed-effects analysis revealed a significant effect of session type, interaction effect of session type x genotype, and interaction effect of session type x genotype x sex (three-way mixed ANOVA; session type: $F_{3,98} = 26.1$, p = 1.1E$^{-11}$; genotype: $F_{1,34} = 0.7$, p = 0.419; sex: $F_{1,34} = 0.004$, p = 0.948; session type x genotype interaction: $F_{3,98} = 3.9$, p = 0.011; session type x sex interaction: $F_{3,98} = 0.9$, p = 0.433; genotype x sex interaction: $F_{1,34} = 3.2$, p = 0.081; session type x genotype x sex interaction: $F_{3,98} = 3.1$, p = 0.028). There was no significant difference between performance of WT female and male mice once they reached the expert performance threshold of d'=2 on the visual task (2.8 ± 0.2 for WT females and 2.7 ± 0.1 for WT males; Mann-Whitney test, p = 0.711; n = 7 WT females and 12 WT males). In addition, there was no significant difference between performance of *Fmr1*$^{-/-}$female and male mice once they reached the expert performance threshold of d'=2 on the visual task (3 ± 0.1 for *Fmr1*$^{-/-}$ females and 3.1 ± 0.3 for *Fmr1*$^{-/-}$ males; Mann-Whitney test, p = 0.837; n = 7 *Fmr1*$^{-/-}$ females and 12 *Fmr1*$^{-/-}$ males). On visual distractor sessions, during trials without distractors, WT female mice performed worse than WT male mice (1.8 ± 0.3 for WT females and 2.5 ± 0.2 for WT males; Mann-Whitney test, p = 0.045). There was no significant difference between performance of *Fmr1*$^{-/-}$ female and male mice (2.3 ± 0.3 for *Fmr1*$^{-/-}$ females and 1.6 ± 0.2 for *Fmr1*$^{-/-}$ males; Mann-Whitney test, p = 0.1). During trials with distractors, there was no significant difference between performance of WT female and male mice (1.7 ± 0.2 for WT females and 2.4 ± 0.2 for WT males; Mann-Whitney test, p = 0.167) and no significant difference between performance of *Fmr1*$^{-/-}$ female and male mice (1.6 ± 0.3 for *Fmr1*$^{-/-}$ females and 1.3 ± 0.3 for *Fmr1*$^{-/-}$ males; Mann-Whitney test, p = 0.606). On the final session of the visual task conducted following the distractor task, there was no significant difference between performance of WT female and male mice (2.5 ± 0.2 for WT females and 2.6 ± 0.1 for WT males; Mann-Whitney test, p = 0.773) and no significant difference between performance of *Fmr1*$^{-/-}$ female and male mice (2.7 ± 0.2 for *Fmr1*$^{-/-}$ females and 2.2 ± 0.3 for *Fmr1*$^{-/-}$ males; Mann-Whitney test, p = 0.181).

In panels A-B, horizontal bars indicate mean and error bars indicate s.e.m; n values are for mice, indicated on each figure. *p < 0.05; **p < 0.01; ***p < 0.001; ****p < 0.0001.

(corresponds to data in Fig. 1 and Supplementary Fig. 2)

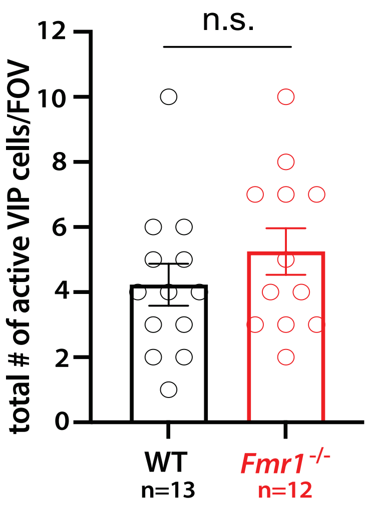


**Supplementary Figure 7:**
There was no significant difference between the total number of active VIP cells per field-of-view (FOV) in WT and *Fmr1*$^{-/-}$ mice (4.2 ± 2.3 for WT mice vs. 5.3 ± 2.5 for *Fmr1*$^{-/-}$ mice; Mann-Whitney test, $p = 0.162$; n = 13 WT mice and 12 *Fmr1*$^{-/-}$ mice).
In panel A, horizontal bars indicate mean and error bars indicate s.e.m; n values are for mice, indicated on the figure. *$p < 0.05$; **$p < 0.01$; ***$p < 0.001$; ****$p < 0.0001$.
(corresponds to data in Fig. 4)

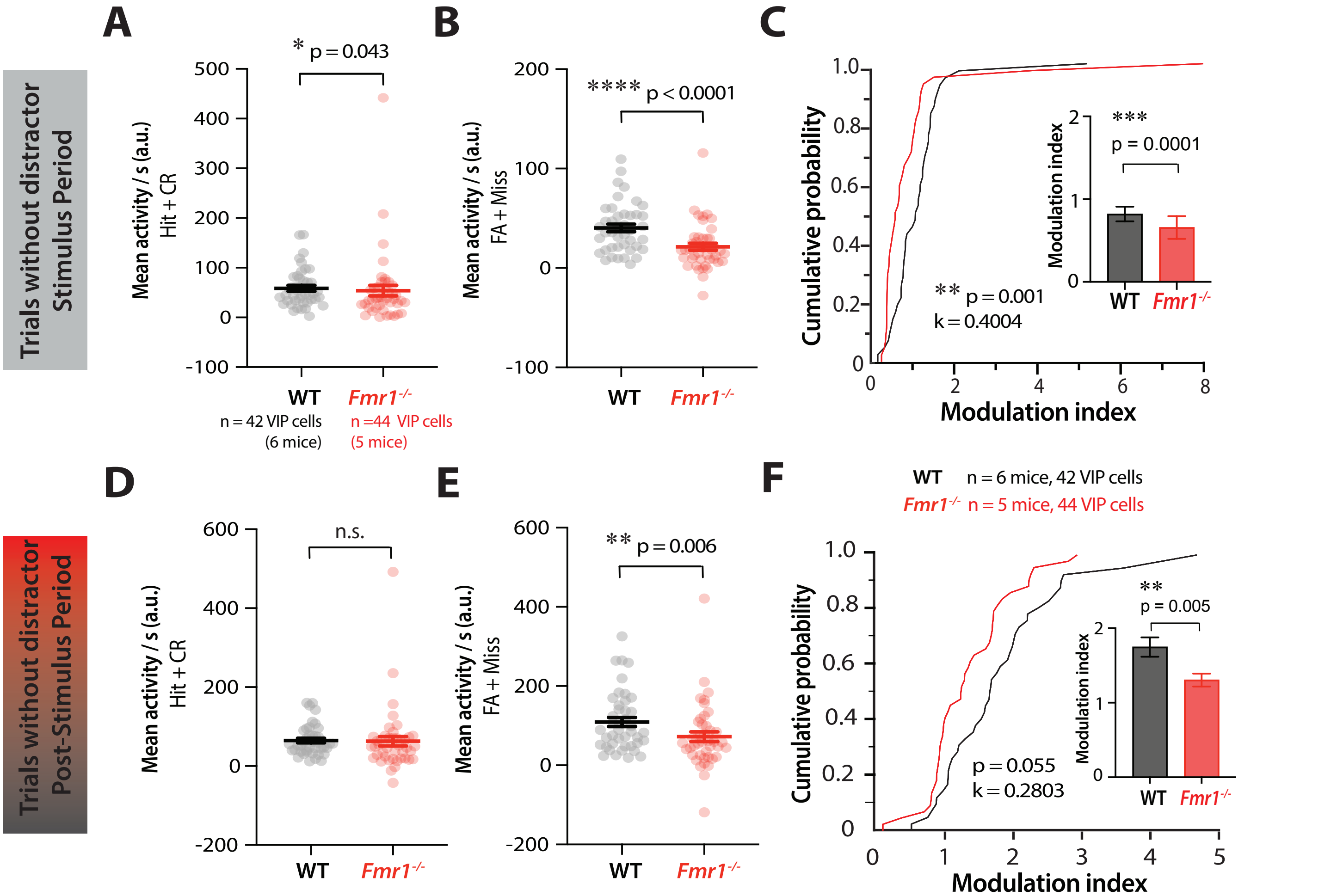


**Supplementary Figure 8:**

**A)** During the stimulus period (0-3 s) of trials without distractors, there was reduced mean VIP activity per second during correct trials for *Fmr1*$^{-/-}$ mice (58.5 ± 38.5 for WT vs. 54 ± 71.4 for *Fmr1*$^{-/-}$; Mann-Whitney test, p = 0.043, Cohen's d = 0.078; n = 6 WT mice, 42 VIP cells and 5 *Fmr1*$^{-/-}$ mice, 44 VIP cells). **B)** There was even greater reduced mean VIP activity per second during error trials for *Fmr1*$^{-/-}$ mice (40.3 ± 25.2 for WT vs. 21.3 ± 23.2 for *Fmr1*$^{-/-}$; Mann-Whitney test, p = 6.3E-5, Cohen's d = 0.784). **C)** A cumulative probability plot showing reduced VIP cell modulation by errors for *Fmr1*$^{-/-}$ mice as measured by the modulation index (two-sample Kolmogorov-Smirnov test, p = 0.001, k = 0.4004). Bar graph inset showing VIP cells were less modulated by errors in *Fmr1*$^{-/-}$ mice (0.8 ± 0.6 for WT vs. 0.7 ± 0.9 for *Fmr1*$^{-/-}$; Mann-Whitney test, p = 0.0001, Cohen's d = 0.214). **D)** During the post-stimulus period (3-6 s for correct trials and 3-12.5 s for error trials) of auditory distractor trials, there was no significant difference in mean VIP activity per second during correct trials (64.8 ± 38.5 for WT vs. 63.0 ± 80.5 for *Fmr1*$^{-/-}$; Mann-Whitney test, p = 0.113).

**E)** There was reduced mean VIP activity per second during error trials for *Fmr1*$^{-/-}$ mice (109.3 ± 75.3 for WT vs. 72.3 ± 80.5 for *Fmr1*$^{-/-}$; Mann-Whitney test, p = 0.006, Cohen's d = 0.475).

**F)** A cumulative probability plot showing VIP cell modulation by errors as measured by the modulation index (two-sample Kolmogorov-Smirnov test, p = 0.055, k = 0.2803). Bar graph inset showing VIP cells were less modulated by errors in *Fmr1*$^{-/-}$ mice (1.7 ± 0.8 for WT vs. 1.3 ± 0.6 for *Fmr1*$^{-/-}$; Mann-Whitney test, p = 0.005, Cohen's d = 0.611). In panels A-F, horizontal bars indicate mean and error bars indicate s.e.m; n values are for mice and cells, indicated on the figure. *p < 0.05; **p < 0.01; ***p < 0.001; ****p < 0.0001.

(corresponds to data in Fig. 5)

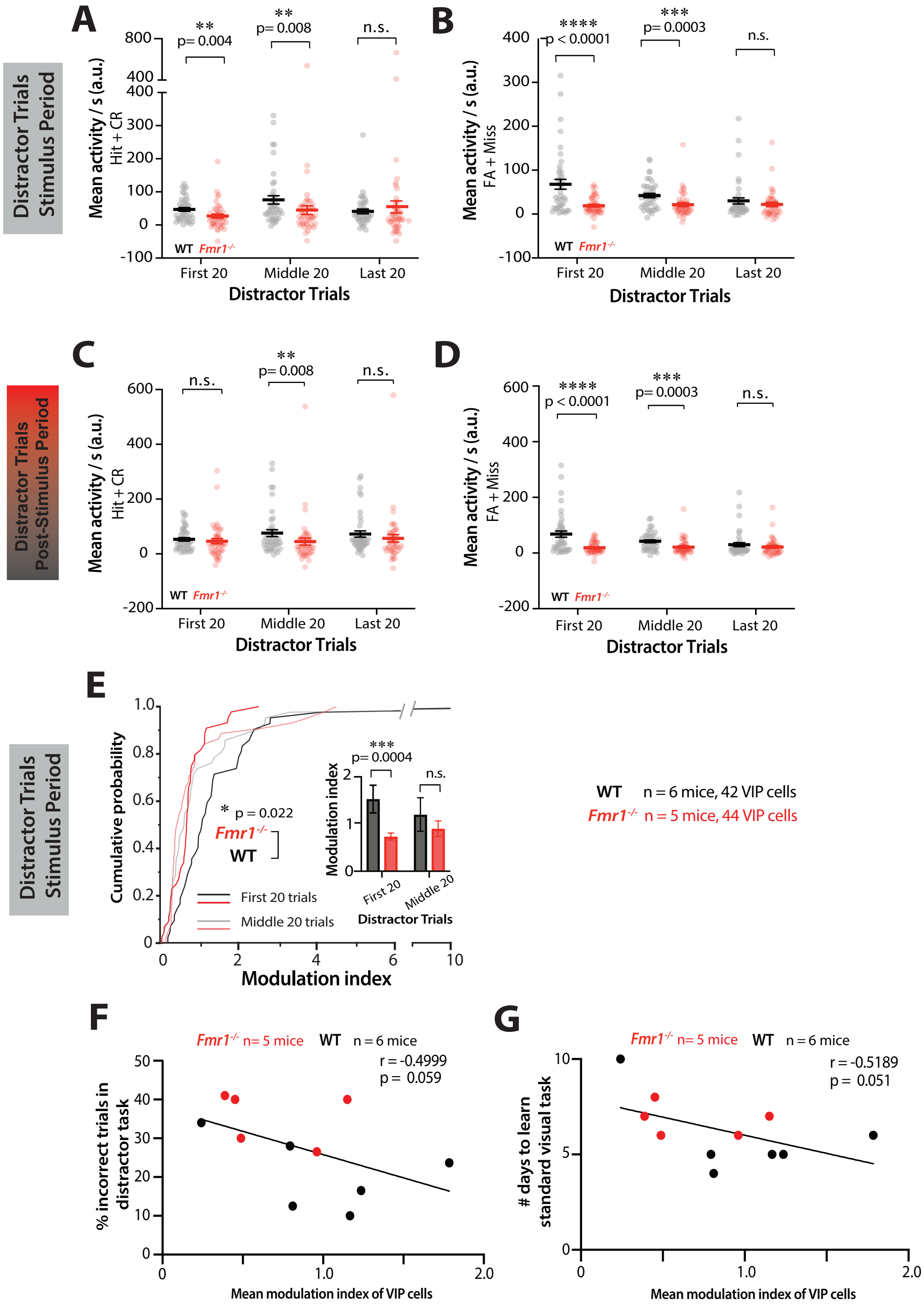


**Supplementary Figure 9**

**Supplementary Figure 9 Legend:**

**A)** During the stimulus period of distractor trials, there was reduced mean VIP activity per second during correct trials for *Fmr1*$^{-/-}$ mice during the first 20 and middle 20 trials (multiple Mann-Whitney tests; first 20: 47.0 ± 34.8 for WT vs. 26.8 ± 38.9 for *Fmr1*$^{-/-}$; p = 0.004, Cohen's d = 0.546; middle 20: 75.9 ± 81.0 for WT vs. 45.0 ± 87.2 for *Fmr1*$^{-/-}$; p = 0.008, Cohen's d = 0.367; last 20: 47.0 ± 34.8 for WT vs. 26.8 ± 38.9 for *Fmr1*$^{-/-}$; p = 0.233; n = 6 WT mice, 42 VIP cells and 5 *Fmr1*$^{-/-}$ mice, 44 VIP cells). There was a significant effect of time (two-way ANOVA; time: $F_{2,168}$ = 3.1, p = 0.048; genotype: $F_{1,84}$ = 1.1, p = 0.297; time x genotype: $F_{2,168}$ = 3, p = 0.054). **B)** During the stimulus period of distractor trials, there was even greater reduced mean VIP activity per second during error trials for *Fmr1*$^{-/-}$ mice during the first 20 and middle 20 trials (multiple Mann-Whitney tests; first 20: 67.9 ± 72.7 for WT vs. 18.9 ± 19.9 for *Fmr1*$^{-/-}$; p = 7.9E-5, Cohen's d = 0.919; middle 20: 42.1 ± 31.9 for WT vs. 21.3 ± 28 for Fmr1-/-; p = 3.3E$^{-4}$, Cohen's d = 0.694; last 20: 30.0 ± 44.9 for WT vs. 21.9 ± 31.9 for *Fmr1*$^{-/-}$; p = 0.403). There was a significant effect of time and effect of genotype (two-way mixed ANOVA; time: $F_{2,164}$ = 5.7, p = 0.004; genotype: $F_{1,84}$ = 15.3, p = 2.0E-4; time x genotype: $F_{2,164}$ = 7.7, p = 0.054). **C)** During the post-stimulus period of distractor trials, there was reduced mean VIP activity per second during correct trials for *Fmr1*$^{-/-}$ mice during the middle 20 trials (multiple Mann-Whitney tests; first 20: 53.2 ± 42.2 for WT vs. 46.4 ± 62.7 for *Fmr1*$^{-/-}$, p = 0.216; middle 20: 72.4 ± 73.2 for WT vs. 56.6 ± 92.4 for *Fmr1*$^{-/-}$, p = 0.008, Cohen's d = 0.19; last 20: 41.7 ± 43.9 for WT vs. 59.8 ± 115.8 for Fmr1-/-; p = 0.183. There was no significant effect or interaction effect (two-way ANOVA; time: $F_{2,168}$ = 1.6, p = 0.202; genotype: $F_{1,84}$ = 1.9, p = 0.171; time x genotype: $F_{2,168}$ = 1, p = 0.358). **D)** During the post-stimulus period of distractor trials, there was reduced mean VIP activity per second during error trials for *Fmr1*$^{-/-}$ mice during the middle 20 trials (multiple Mann-Whitney tests; first 20: 154.0 ± 128.5 for WT vs. 56.8 ± 78.4 for *Fmr1*$^{-/-}$, p = 7.9E-5, Cohen's d = 0.913 ; middle 20: 119.0 ± 87.0 for WT vs. 62.9 ± 98.3 for *Fmr1*$^{-/-}$, p = 3.3E-4, Cohen's d = 0.605; last 20: 59.2 ± 57.6 for WT vs. 79.0 ± 116.7 for *Fmr1*$^{-/-}$, p = 0.403. There was a significant effect of time, effect of genotype, and time x genotype interaction effect (two-way mixed ANOVA; time: $F_{2,164}$ = 5.7, p = 0.004; genotype: $F_{1,84}$ = 15.3, p = 2.0E-4; time x genotype: $F_{2,164}$ = 7.7, p = 7.0E-4). **E)** A cumulative probability plot showing reduced VIP cell modulation by errors for *Fmr1*$^{-/-}$ mice during the stimulus period on the first 20 distractor trials (two-sample Kolmogorov-Smirnov test, p = 3.1E-4, k = 0.438) and no significant difference on the middle 20 distractor trials (two-sample Kolmogorov-Smirnov test, p = 0.365, k = 0.193). Bar graph inset showing VIP cells were less modulated by errors in *Fmr1*$^{-/-}$ mice on the first 20 distractor trials (1.5 ± 1.9 for WT vs. 0.7 ± 0.5 for *Fmr1*$^{-/-}$, Mann-Whitney test, p = 4.30E-4, Cohen's d = 0.569) and there was no significant difference on the middle 20 distractor trials (1.2 ± 2.3 for WT vs. 0.9 ± 1.1 for *Fmr1*$^{-/-}$; Mann-Whitney test, p = 0.494). There was a significant effect of genotype (two-way ANOVA; time: $F_{1,84}$ = 0.1, p = 0.748; genotype: $F_{1,84}$ = 5.4, p = 0.022; time x genotype: $F_{1,80}$ = 0.9, p = 0.33).

**F)** The modulation index of VIP cells during the stimulus period of all auditory distractor trials was averaged per mouse and was trending towards a negative correlation with percent incorrect (%) on the distractor task (Pearson's r, r = -0.4999, p = 0.059). A regression line is fitted to the data points. **G)** The modulation index of VIP cells during the stimulus period of all auditory distractor trials was trending towards a negative correlation with number of days to learn the standard visual task (Pearson's r, r = -0.5189, p = 0.051).

In panels A-E, horizontal bars indicate mean and error bars indicate s.e.m; n values are for mice and cells, indicated on the figure. *p < 0.05; **p < 0.01; ***p < 0.001; ****p < 0.0001.

(corresponds to data in Fig. 5)
